# A conserved mechanism of LRRC8 channel inhibition by distinct drugs

**DOI:** 10.1101/2025.02.24.639894

**Authors:** Toshiki Yamada, Paola Bisignano, Erkan Karakas, Jerod S. Denton

## Abstract

We employed a Leucine Rich Repeat Containing 8 (LRRC8) channel chimera, termed 8C-8A(IL1^25^), to investigate the molecular mechanism of action of the novel volume-sensitive anion channel (VRAC) inhibitor, zafirlukast. 8C-8A(IL1^25^) comprises LRRC8C (8C) and 25 residues from LRRC8A (8A) intracellular loop 1 (IL1) and forms volume-sensitive, structurally defined heptameric channels with normal pharmacological properties. In silico docking and modeling with AlphaFold3 identified a putative zafirlukast binding site comprising the amino (N)-terminal domain (NTD) and inter-subunit fenestrae between transmembrane (TM) helices 1 and 2. Consistent with this model, mutations in NTD, TM1, and TM2 alter 8C-8A(IL1^25^) and heteromeric 8A/8C sensitivity to zafirlukast and the structurally distinct drug pranlukast. Inhibition is not mediated by extracellular pore block or the so-called lipid gate. Mutations or low pH conditions that enhance voltage-dependent inactivation also increase zafirlukast sensitivity. We propose zafirlukast and pranlukast promote channel inactivation through destabilization of the pore.

## Introduction

Volume-Regulated Anion Channels (VRACs) participate in essential physiological processes, such as cellular osmoregulation, cell migration, and cell death^1–4^. Since the identification of the Leucine Rich Repeat Containing 8 (LRRC8) gene family as the primary component of VRACs in 2014^5, 6^, the use of genetic techniques has uncovered new and potentially “druggable” roles in cardiovascular, immunological, metabolic, and neurological disorders ^7–15^. However, exploring the therapeutic potential of LRRC8/VRACs in cell, tissue, and animal disease models is hampered by the lack of pharmacological tools for selectively modulating channel activity. The current best-in-class LRRC8/VRAC inhibitor in terms of potency and selectivity within the anion channel superfamily is 4-[(2-Butyl-6,7-dichloro-2-cyclopentyl-2,3-dihydro-1-oxo-1H-inden-5-yl)oxy]butanoic acid (DCPIB). However, even DCPIB suffers from moderate potency and extensive off-target activity^16–20^, highlighting the need for better tool compounds^21^.

To expand the toolkit for probing the druggability of LRRC8/VRAC, we previously screened a library of 1,184 FDA-approved drugs for novel activators and inhibitors of endogenous VRACs expressed in HEK-293 cells. One novel VRAC inhibitor identified in the screen is pranlukast (Onon^®^)^22^, a nanomolar-affinity antagonist of the cysteinyl leukotriene receptor-1 (CysLT1R) (Fig 1). Pranlukast is widely prescribed in Japanese markets to treat airway constriction associated with allergic rhinitis and asthma^23^. In both conditions, inflammatory leukotrienes released from immune cells activate CysLT1R expressed on airway smooth muscle cells, triggering Gq-coupled calcium (Ca^2+^) signaling, myocyte contraction and airway constriction. Pranlukast inhibits pro-inflammatory signaling by displacing the leukotriene molecule from its binding site in CysLT1R.^24^

**Figure 1.**
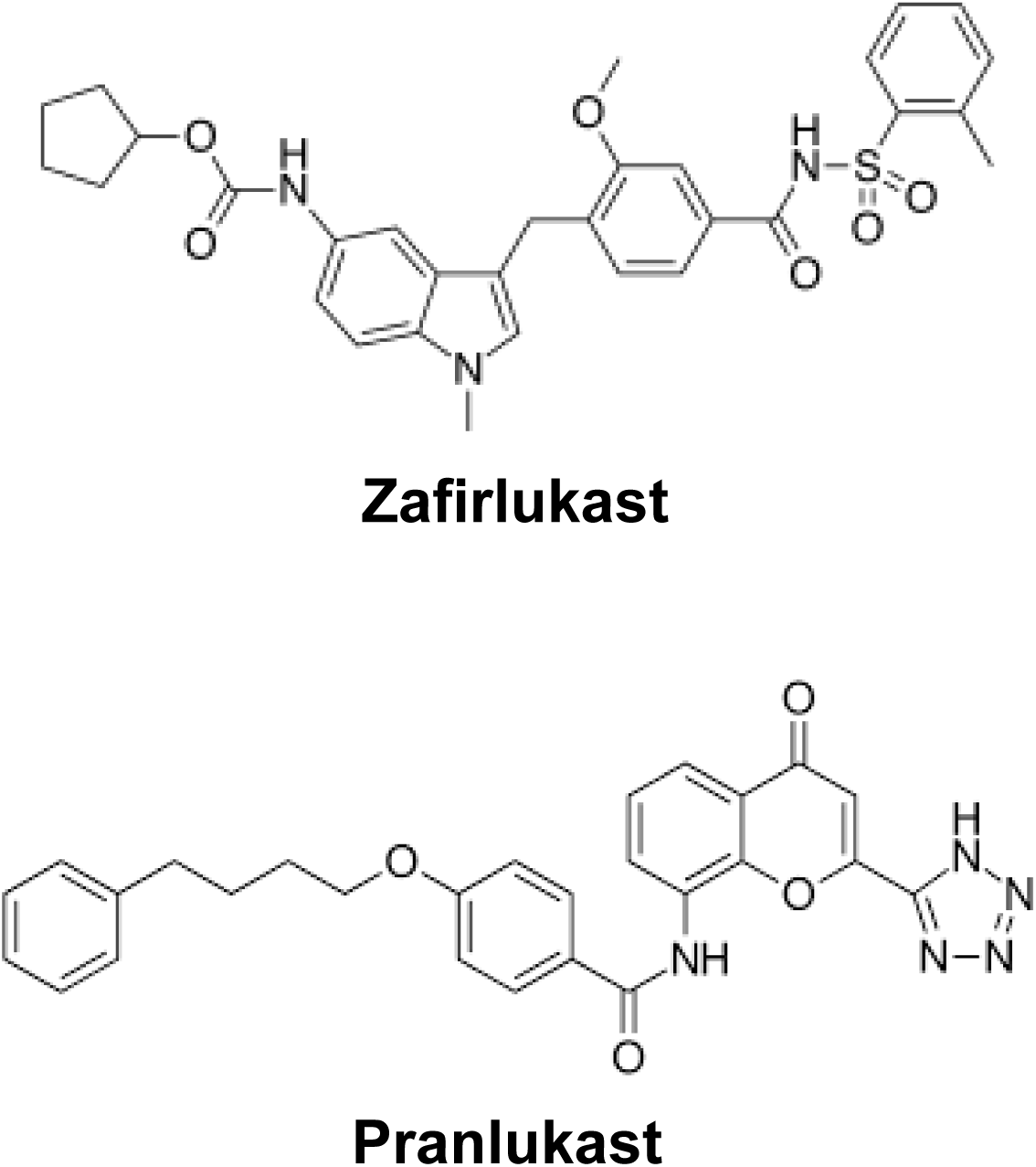
Chemical structures of zafirlukast and pranlukast.

In voltage clamp experiments, pranlukast rapidly (time constant, τ, ∼1 min), reversibly and dose-dependently inhibited swelling-activated VRAC currents at single micromolar (µM) doses and a maximal efficacy (E_max_) of ∼50%. Additionally, and somewhat surprisingly, we found that the structurally distinct CysLT1R antagonist, zafirlukast (Accolate®) (Fig. 1), inhibits VRAC with similar potency but with ∼100% E_max_. The discovery of pranlukast and zafirlukast as VRAC inhibitors suggested that the drugs inhibit channel activity indirectly by interfering with CysLT1R signaling. However, several lines of evidence indicated that channel inhibition occurs independently of CysLT1R-dependent signaling, suggesting the drugs act directly on VRACs^21, 22^.

The LRRC8 family consists of 5 genes: *LRRC8A*, *LRRC8B*, *LRRC8C*, *LRRC8D*, and *LRRC8E*. The formation of volume-sensitive anion channels with native biophysical properties requires the assembly of the essential subunit, LRRC8A, with LRRC8C, LRRC8D, or LRRC8E, or a combination thereof^5^. Each LRRC8 subunit consists of an intracellular amino (N)-terminal domain, a transmembrane (TM) domain composed of 4 TM helices, two extracellular loop domains, an intracellular loop domain, and intracellular Leucine-Rich-Repeat (LRR) domains. A growing number of cryo-EM structures of homomeric and heteromeric LRRC8 channel structures has established that LRRC8s belong to the so-called “large pore” family of membrane channels, which also includes pannexins, connexins, and CALHMs^11, 25–30^. Some structures have revealed lipids that penetrate through the channel wall and occlude the ion conduction pathway^26, 31–33^. Charged mutations at the outer pore mouth cause electrostatic repulsion of lipids from the pore and constitutive channel activation, supporting the notion that pore-resident lipids play a direct role in LRRC8 channel gating^31^.

Investigating the mechanism of drug action on LRRC8/VRACs is complicated by the heteromeric nature of the channel. The uncontrollable, indeterminate, and variable subunit stoichiometry and position give rise to structurally heterogenous populations of LRRC8 channels, rendering the molecular context of introduced mutations different between subpopulations of channels. We recently developed a series of LRRC8 chimeric constructs that can be functionally expressed as homomeric channels and thereby circumvent many complications of heteromeric channels^34^. One chimera, termed 8C-8A(IL1^25^), comprises the LRRC8C protein in which 25 amino acids from the LRRC8A intracellular loop 1 (IL1) are swapped into the corresponding region of LRRC8C. 8C-8A(IL1^25^) exhibits native biophysical and regulatory properties and sensitivity to DCPIB^34, 35^. Cryo-EM analysis indicates that 8C-8A(IL1^25^) forms large-pore heptameric channels^26^ whose overall structure is very similar to that of LRRC8A/LRRC8C heteromeric channels, which are hexamers^27, 31, 32^.

In the present study, we employed the 8C-8A(IL1^25^) chimera as a structurally defined homomeric channel model to study the molecular mechanism of action of zafirlukast. Attempts to generate a co-structure of zafirlukast with 8C-8A(IL1^25^) chimera were unsuccessful, so we turned to computational techniques. In silico docking against the 8C-8A(IL1^25^) cryo-EM structure and AlphaFold modeling led to the identification of a putative zafirlukast binding site located between the N-terminal domain (NTD) and TM helices 1 (TM1) and 2 (TM2) near inter-subunit fenestrae believed to interact closely with plasma membrane phospholipids. Consistent with this model, mutations in the NTD, TM1, and TM2 lead to striking changes to zafirlukast and pranlukast sensitivity of 8C-8A(IL1^25^) as well as LRRC8A/LRRC8C heteromeric channel. Additional evidence suggests that drug inhibition is not mediated by the closure of the “lipid gate” but rather by constricting or collapsing the pore. This may represent a general mechanism of LRRC8/VRAC channel inhibition by lipophilic drugs.

## Methods

### Drugs and other reagents

Zafirlukast and pranlukast were purchased from Tocris or Cayman as dry powders, dissolved as 100 and 50 mM DMSO stock solutions, respectively, and diluted in experimental buffers immediately before use. MTSET reagent was purchased from Toronto Research Chemicals, dissolved as a 400 mM stock solution in water, stored at -80°C in single-use aliquots, and diluted in bath solution immediately before application.

### Molecular biology

Site-directed mutagenesis was performed using Phusion High-Fidelity PCR Master Mix (ThermoFisher Scientific) according to the manufacturer’s instructions. The open reading frame was fully sequenced to ensure the fidelity of mutagenesis.

### Cell lines and transfections

HCT116 cells in which all five *LRRC8* genes have been disrupted using CRISPR-Cas9 (HCT*^LRRC8-/-^*) were a gift from Dr. Thomas Jentsch (Leibniz-Forschungsinstitut für Molekulare Pharmakologie (FMP), Berlin Germany). Cells were transfected using Turbofection 8.0 Transfection Reagent (OriGene Technologies) with 0.125 μg GFP and 0.125 to 1 μg various LRRC8 plasmids according to the manufacturer’s instructions. Transfected cells were studied 36-48 hours after transfection.

### Whole-cell patch clamp electrophysiology

Whole-cell voltage-clamp electrophysiology was performed essentially as described previously (Yamada et al, 2020). Briefly, transfected cells were identified by their GFP fluorescence and patch clamped using a bath solution containing 75 mM CsCl, 5 mM MgSO_4_, 2 mM Ca gluconate, 12 mM HEPES, 8 mM Tris, 5 mM glucose, 2 mM glutamine, and 115 mM sucrose, pH 7.4 adjusted with CsOH (300 mOsm), and a pipette solution containing 126 mM CsCl, 2 mM MgSO_4_, 20 mM HEPES, 1 mM EGTA, 2 mM Na-ATP, 0.5 mM Na-GTP and 10 mM sucrose, pH 7.2 adjusted with CsOH (275 mOsm). Low intracellular ionic strength pipette solution was prepared by reducing the CsCl concentration from 126 mM (normal ionic strength) to 26 mM (low ionic strength). Cells were swollen by exposure to hypotonic bath solution (250 mOsm). Osmolality was lowered by removing sucrose. All experiments were performed at room temperature.

Drug concentration-response relationships were established in voltage-ramp experiments, as follows. LRRC8/VRAC currents were evoked from a holding potential of -30 mV by first stepping to -100 mV for 15 msec and then ramping the voltage to +100 mV over 1 sec. The membrane potential was subsequently stepped back to -30 mV for 4 sec until repeating the step-ramp protocol. Voltage-step protocols were also used to characterize the voltage-dependent properties of LRRC8/VRAC currents. The membrane potential was first stepped to -80 mV for 500 msec and then stepped for 2 sec from -120 to +120 mV in 20 mV increments. The instantaneous current was used for plotting current-voltage relationships. Rectification rates were calculated as the absolute value of the maximum current at +120 mV divided by that at -120 mV. Inactivation ratios are relative values of the steady-state current divided by the instantaneous current at +120 mV. Time constants were estimated by fitting a single exponential function to current at +120 mV for 2 sec using Clampfit 10 (Molecular devices). For time constants of zafirlukast inhibition, the tau values and statistical significance were determined by Plateau followed by one phase decay equation on GraphPad Prism 10 software.

### Docking Methods

The binding mode of zafirlukast to 8C-8A(IL1^25^) was predicted using PDB ID 8DXN. All computational studies were carried out with the *Molecular Operating Environment* (MOE) suite, version *2022.02*^36^. Initially, the protein structure and the zafirlukast molecule were prepared in a format ready for docking. The *QuickPrep* and *Wash* utilities in MOE were utilized to assign bond orders and protonation states at pH 7.4, optimize intramolecular hydrogen bonds, remove salts and solvents, and enumerate the ligand to generate all possible tautomers. Subsequently, the *SiteFinder* algorithm was used to screen all possible druggable sites within the protein structure. Dummy atoms were then generated at the top ranked sites identified by SiteFinder, serving as reference points for the pose search during the subsequent docking procedure. Binding pose predictions were performed using the general docking procedure, increasing the pose search and refinement to 30 poses and permitting the relaxation of protein residues within one solvation shell of the bound ligand pose. The selection of docking poses was based on visual inspection and docking scores.

### Modeling of the NTB with AlphaFold 3

The NTD comprising residues 1 to 15 is not resolved in 8C-8A(IL1^25^) chimera structures. To model the NTD, we used the Alpha Fold 3 server^37^. The prediction was performed using the 8C-8A(IL1^25^) chimera sequence, which includes residues 1 to 411 (excluding the LRR domains), and searching for a heptameric structure. One of the subunits from the predicted structure was superimposed onto each subunit of the 8C-8A(IL1^25^) chimera (PDB ID: 8DXN) to obtain a predicted structure of 8C-8A(IL1^25^) chimera with the NTD.

### Statistical analysis

Data are presented as mean ± standard error of the mean (SEM), n represents the number of cells in patch clamp recordings. GraphPad Prism 10 software was used for all statistical analysis. Concentration response curves were generated by fitting of four parameters Hill equation with constraints bottom and top to 0 and 100, respectively. Statistical significance was determined by Student’s unpaired or paired t-test for two groups, one-way analysis of variance (ANOVA) followed by Dunnett’s multiple comparisons test for groups greater than two.

## Results

### Characteristics of 8C-8A(IL1^25^) current inhibition by zafirlukast

The inhibitory effects of zafirlukast on swelling-activated 8C-8A(IL1^25^) currents were evaluated in transfected HCT *^LRRC8-/-^* cells patch clamped with normal ionic strength pipette solution (i.e., 126 mM CsCl). As reported previously^34^, 8C-8A(IL1^25^) channels exhibit low basal activity in isotonic bath (data not shown) but undergo striking activation by hypotonic cell swelling (Fig. 2a, left panel). Swelling-activated currents are outwardly rectifying and undergo voltage-dependent inactivation (VDI) at large positive test potentials, similar to native VRAC currents^38–40^. Subsequent bath application of 30 µM zafirlukast in hypotonic buffer led to a complete inhibition of 8C-8A(IL1^25^) current across all voltages tested (Fig. 2b, middle panel). Drug inhibition occurred with a time constant, tau, of 24.4 ± 1.1 sec (+100 mV) and 27.2 ± 1.2 sec (-100 mV) and was not reversible during a 5-min washout period (Fig. 2a-b). Dose-response experiments revealed that zafirlukast inhibits 8C-8A(IL1^25^) in a voltage-independent manner with IC_50_ values of 8.7 (+100 mV) and 9.3 µM (-100 mV) (Fig. 2c and Table 1). We also evaluated the characteristics of zafirlukast inhibition of 8C-8A(IL1^25^) currents activated with low intracellular ionic strength (26 mM CsCl) (Fig. 2d). Under these conditions, zafirlukast inhibited 8C-8A(IL1^25^) with a tau of 21.2 ± 1.0 sec (+100 mV) and 22.9 ± 1.3 sec (-100 mV) (Fig. 2e) and IC_50_ values of 8.7 µM (+100 mV) and 9.8 µM (-100 mV) (Fig. 2f), which are not statistically different from those recorded under normal intracellular ionic strength conditions (Table 1). One notable difference, however, was that zafirlukast inhibition is partially reversible under low intracellular ionic strength conditions (Figs. 2e).

**Figure 2.**
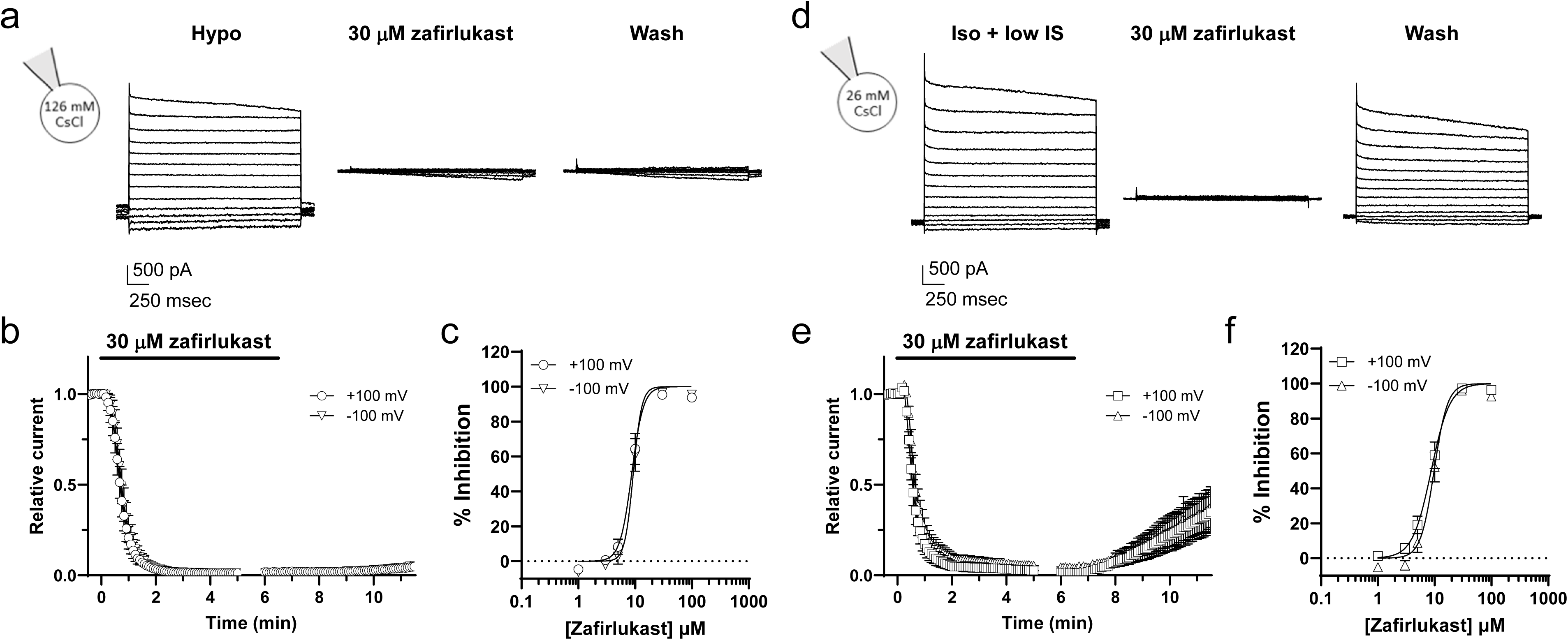
Characteristics of zafirlukast-dependent inhibition of 8C-8A(IL1^25^) currents. **a)** Representative whole-cell 8C-8A(IL1^25^) currents recorded from a cell dialyzed with normal intracellular ionic strength (126 mM CsCl) pipette solution cell swelling with hypotonic buffer (left panel), after bath application of 30 µM zafirlukast (middle panel), and following washout of the drug (right panel). **b)** Time-course of zafirlukast inhibition of swelling-activated 8C-8A(IL1^25^) currents and drug washout. Data are mean ± SEM (n=10) current at +100 mV and -100 mV normalized to that measured before drug addition. **c)** Mean ± SEM (**n=4-8**) CRC of zafirlukast inhibition at +100 mV and -100 mV. **d)** Representative 8C-8A(IL1^25^) currents recorded from a cell dialyzed with low intracellular ionic strength (26 mM CsCl) pipette solution in isotonic buffer (left panel), after bath application of 30 µM zafirlukast (middle panel), and following washout of the drug (right panel). **e)** Time-course of zafirlukast inhibition and drug washout. Data are mean ± SEM (n=4) currents at +100 mV and -100 mV. **f)** Mean ± SEM (n=7) CRC of zafirlukast inhibition at +100 mV and -100 mV.

**Table 1.** Zafirlukast sensitivity of LRRC8 constructs.

| Construct | IC<sub>50</sub> (μM) |  | Hill coefficient |  |
| --- | --- | --- | --- | --- |
|  | +100 mV | -100 mV | +100 mV | -100 mV |
| **Chimera** |  |  |  |  |
| **8C-8A(IL1<sup>25</sup>)** | 8.7 | 9.3 | 4.22 | 5.81 |
| **8C-8A(IL1<sup>25</sup>)** | 8.7 | 9.8 | 2.64 | 3.67 |
| **Low ionic strength** |  |  |  |  |
| **8C-8A(IL1<sup>25</sup>)-E6A** | 0.47 | 0.56 | 0.87 | 0.96 |
|  | P<0.0001 | P<0.0001 | P<0.0001 | P<0.0001 |
| **8C-8A(IL1<sup>25</sup>)-C43G** | 2.6 | 2.7 | 1.80 | 1.90 |
|  | P<0.0001 | P<0.0001 | P<0.005 | P<0.005 |
| **8C-8A(IL1<sup>25</sup>)-Q9C/C43G** | 9.8 | 10.0 | 2.31 | 2.38 |
|  | P<0.0001* | P<0.0001* |  |  |
| **8C-8A(IL1<sup>25</sup>)-S11C/C43G** | 12.6 | 11.6 | 1.56 | 1.57 |
|  | P<0.005 |  | P<0.001 | P<0.0005 |
|  | P<0.0001* | P<0.0001* | P<0.0001* |  |
| **8C-8A(IL1<sup>25</sup>)-M48D** | 23.0 | 17.0 | 0.78 | 0.92 |
|  | P<0.0001 | P<0.0001 | P<0.0001 | P<0.0001 |
| **8C-8A(IL1<sup>25</sup>)-L105R** | 6.1 | 5.8 | 2.19 | 2.60 |
|  | P<0.005 | P<0.005 | P<0.01 | P<0.01 |
| **8C-8A(IL1<sup>25</sup>)-K125Q** | 4.2 | 4.2 | 1.69 | 1.80 |
|  | P<0.0001 | P<0.0001 | P<0.001 | P<0.001 |
| **8C-8A(IL1<sup>25</sup>)-Y126F** | 0.79 | 0.93 | 0.90 | 1.04 |
|  | P<0.0001 | P<0.0001 | P<0.0001 | P<0.0001 |
| **8C-8A(IL1<sup>25</sup>)-I133F** | 5.8 | 6.9 | 1.52 | 1.63 |
|  | P<0.005 | P<0.05 | P<0.0001 | P<0.0005 |
| **8C-8A(IL1<sup>25</sup>)-L136K** | 0.088 | 0.099 | 0.66 | 0.81 |
|  | P<0.0001 | P<0.0001 | P<0.0001 | P<0.0001 |
| **Heteromer** |  |  |  |  |
| **8A/8C** | 17.6 | 15.2 | 1.34 | 1.67 |
| **8A-E6A/8C E6A** | 1.6 | 1.6 | 1.39 | 1.23 |
|  | P<0.0001 | P<0.0001 |  |  |
| **8A-T48D/8C-M48D** | 45.5 | 56.0 | 1.23 | 1.53 |
|  | P<0.0001 |  |  |  |
| **8A-124F/8C-Y126F** | 19.6 | 18.5 | 1.53 | 1.78 |
| **8A-L134K/8C-L136K** | 3.4 | 3.9 | 0.99 | 1.01 |
|  | P<0.0001 | P<0.0001 |  | P<0.05 |

P values reflect comparisons of wild type 8C-8A(IL1<sup>25</sup>) or 8A/8C with mutants. Asterisks indicate comparisons with 8C-8A(IL1<sup>25</sup>)-C43G.

The sidedness of zafirlukast inhibition was evaluated by applying the drug extracellularly in the bath solution or intracellularly via the pipette solution. In bath application experiments, 8C-8A(IL1^25^) channels were first closed with hypertonic shrinkage for 6 min to ensure that zafirlukast cannot permeate the pore and accumulate intracellularly. Cells were then exposed to either DMSO (solvent control) or 30 µM zafirlukast for 5 min in the continued presence of hypertonic bath, washed continuously for 3.5 min with hypertonic buffer, and then subjected to hypotonic cell swelling for 6 min to activate 8C-8A(IL1^25^). Swelling-induced currents in DMSO-treated cells were significantly larger than those treated with zafirlukast (Fig. 3a-b), indicating that not only can zafirlukast access its binding site from the extracellular side of the channel, but that the binding site is accessible in fully closed channels. We next tested if zafirlukast can inhibit 8C-8A(IL1^25^) when applied to the intracellular side of the channel by delivering the drug directly through the pipette solution. As shown in Fig. 3c-d, intracellular delivery of 30 µM zafirlukast through the pipette solution did not prevent 8C-8A(IL1^25^) current activation with cell swelling or subsequent inhibition by zafirlukast.

**Figure 3.**
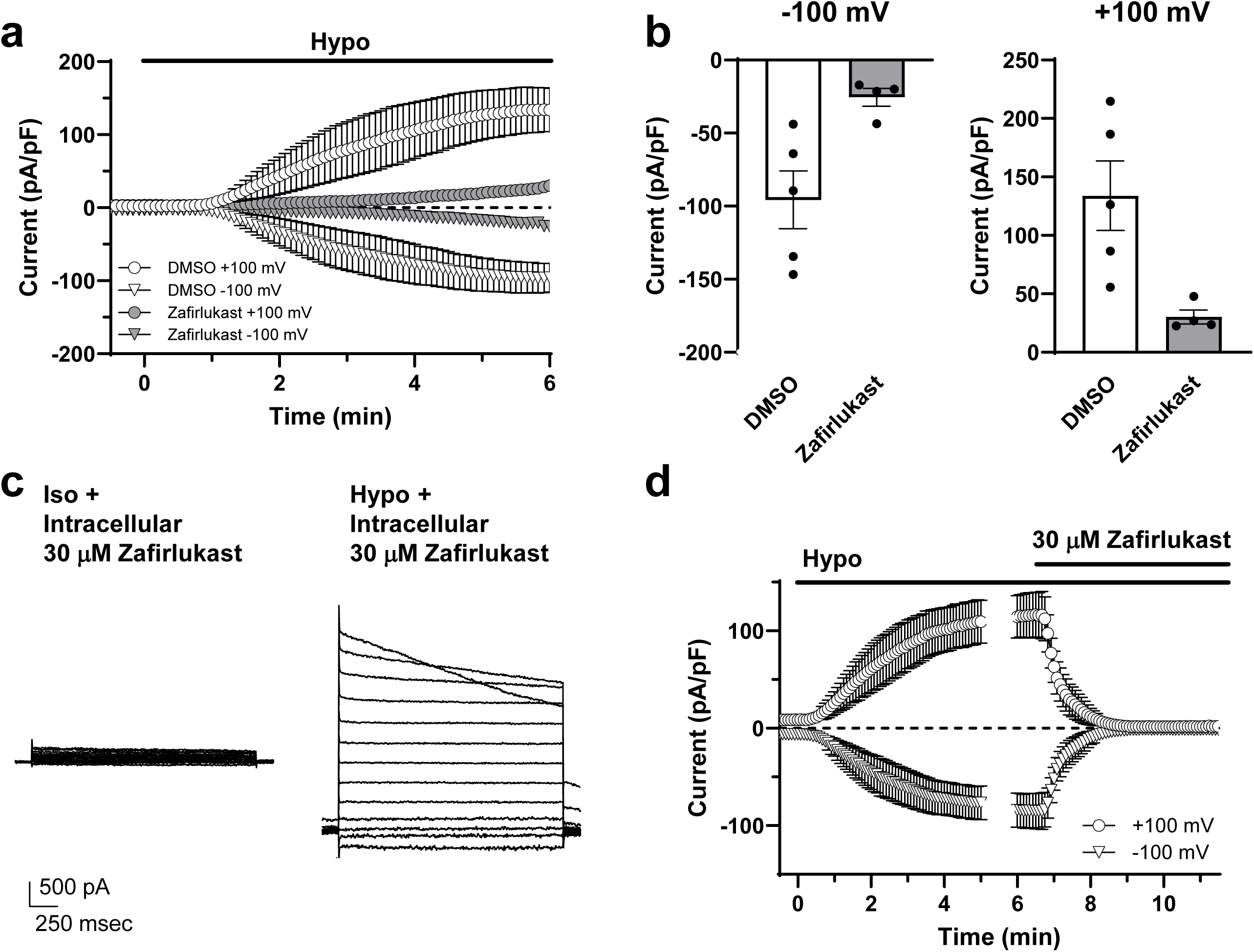
Sidedness of zafirlukast action. **a)** Inhibition of 8C-8A(IL1^25^) channels in the closed state. Cells were shrunken with hypertonic saline for 6 min before applying DMSO (solvent control) or 30 µM zafirlukast in hypertonic buffer before washout and subsequent activation of the channel with hypotonic swelling. Data are means ± SEM current density (n=4-5). **b)** Mean ± SEM steady-state swelling-activated current density recorded at -100 mV (left panel) and +100 mV (right panel)(n=4-5). **c)** No effect of intracellular zafirlukast on 8C-8A(IL1^25^) channels. Representative 8C-8A(IL1^25^) currents evoked from a cell dialyzed with 30 µM zafirlukast before (left panel) and after (right panel) hypotonic swelling. **d)** Mean ± SEM current density at +100 mV (circles) or -100 mV (inverted triangles) vs time plots (n=4).

### Computational docking of zafirlukast into the 8C-8A(IL1^25^) structure

Attempts to solve the co-structure of zafirlukast bound to 8C-8A(IL1^25^) chimera were unsuccessful, so we turned to computational methods to identify potential binding sites. The computational studies were performed using the MOE suite to identify the putative binding site of zafirlukast at the 8C-8A(IL1^25^) chimera. Druggable areas identified using the Site Finder tool revealed the subunit interfaces within the TMs as the primary sites for zafirlukast binding. Given the high similarity of the interfaces within the homomeric heptamers, we focused the docking studies on a single interface. A broad pose search was conducted for zafirlukast, followed by induced fit refinement, allowing relaxation of protein residues within 6 Å of the predicted ligand pose. The combined analysis of rigid docking and induced fit refinement resulted in 18 subclusters and six singletons, three from each method. After visual inspection, the most populated binding modes were selected based on consensus between rigid docking and induced fit poses. The docking analysis reveals that the zafirlukast binding site is buried at the subunit interface within the TM domain (Fig. 4). Part of the docked zafirlukast molecules faces toward the pore but does not appear to affect solvent-accessible pore dimensions (Fig. 4c-e). The other side of the docked molecules faces towards the hydrophobic core of the membrane bilayer. Notably, this putative binding site is occupied by acyl chains of phospholipids in the cryo-EM structures of 8C-8A(IL1^25^) chimera^26^. Considering their hydrophobic nature, zafirlukast molecules may plausibly replace the lipid molecules at the subunit interfaces. The key residues that are at a close distance to the docked molecules are C43, M48, I133, and L136. No docking poses were identified near L105, which forms the tightest constriction at the ECD. *Mutagenesis analysis of the zafirlukast docking model in 8C-8A(IL1^25^)*

**Figure 4.**
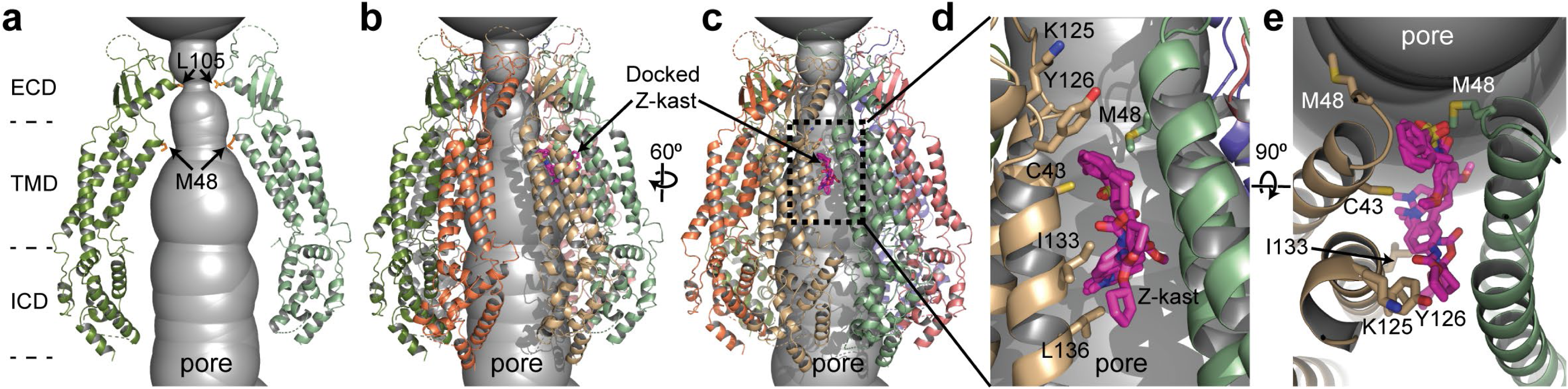
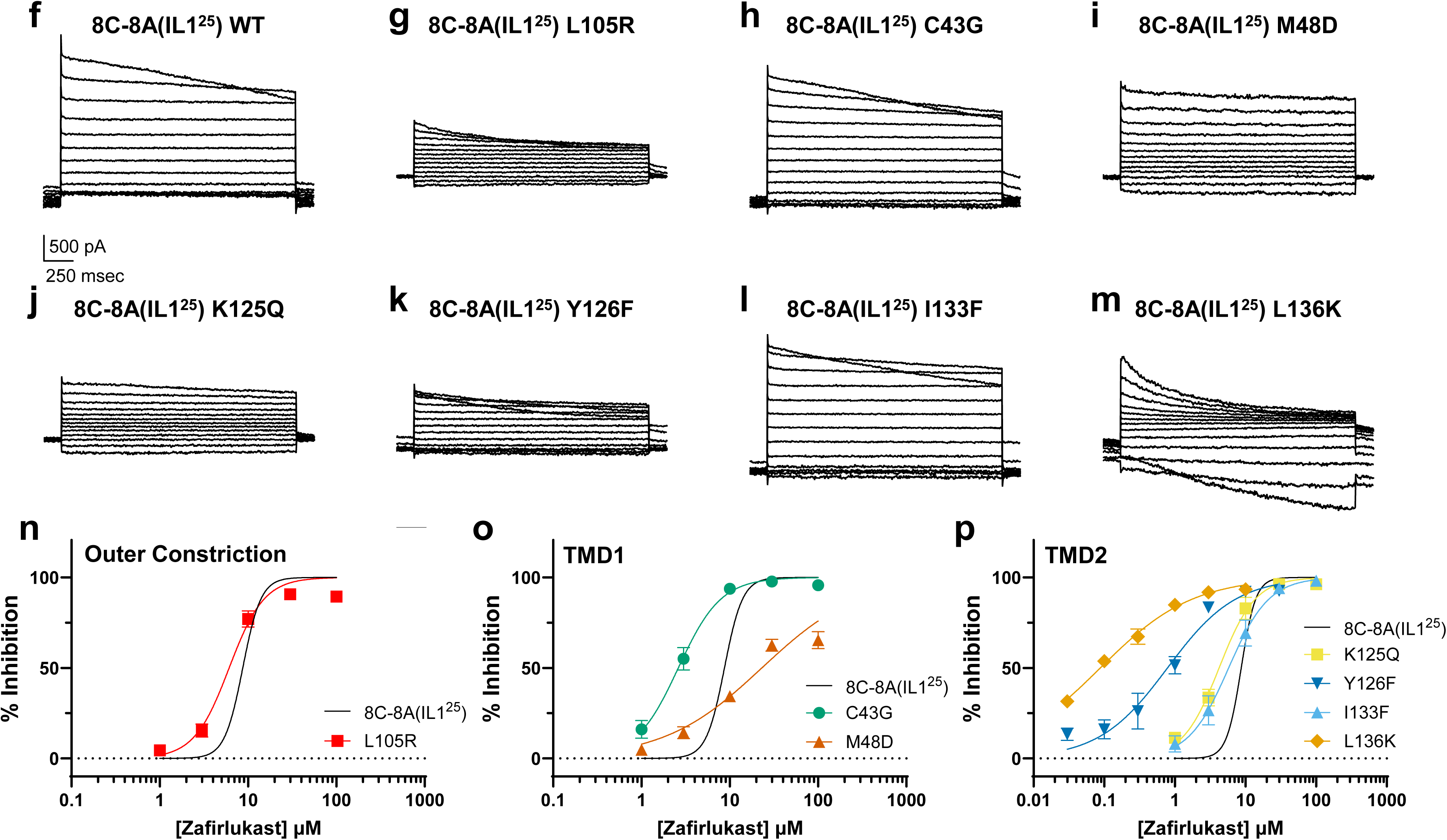
Analysis of docking site mutations on 8C-8A(IL1^25^) sensitivity to zafirlukast. **a-e)** Cryo-EM structure of 8C-8A(IL1^25^) (PDB ID:8DXN) along with the docked zafirlukast molecules (magenta) and the solvent-accessible permeation pathway (gray surface) calculated using HOLE^54^. **a)** Two opposing subunits are shown to highlight the positions of M48 and L105. **b-c)** Two different views of the entire pore region of 8C-8A(IL1^25^) and docked zafirlukast molecules at one of the subunit interfaces. Only the top five docking poses for zafirlukast are shown. **d-e)** Close-up view of the putative binding pocket for zafirlukast from two angles. Residues used for mutational analysis are shown as sticks. **f-m)** Representative whole-cell currents from the indicated mutants**. n)** CRC data from outer constriction mutant 8C-8A(IL1^25^)-L105R. Fit of WT 8C-8A(IL1^25^) CRC data (black line) from Fig. 2c is shown for comparison. **o)** CRC data for WT 8C-8A(IL1^25^) (black line), and TM1 mutants C43G and M48D. p) CRC data for WT 8C-8A(IL1^25^) (black line) and TM2 mutants K125Q, Y126F, I133F, and L136K. Data are means ± SEM (WT n = 4-8, L105 n = 5-6, C43G n = 5-6, M48D n = 4-6, K125Q, n = 5, Y126F n = 5-7, I133F n = 5, L136K n = 5-7).

The narrowest part of the resolved 8C-8A(IL1^25^) channel pore is formed by leucine 105 (L105) (Fig. 4a)^26^, which is equivalent to arginine 103 (R103) in LRRC8A. 8C-8A(IL1^25^)-L105 residue was of interest because R103 forms the binding site for DCPIB in homohexameric 8A channels^33, 35^. Despite our docking analysis failing to identify a stable binding pose of zafirlukast near L105, we evaluated the drug sensitivity of 8C-8A(IL1^25^)-L105R channels, as an initial validation of our model. 8C-8A(IL1^25^)-L105R channels exhibited normal volume sensitivity (data not shown), reduced rectification (Table 2), enhanced VDI (Fig. 4g and Table 2), and slightly lower zafirlukast IC_50_ (Fig. 4n and Table 1). Thus, consistent with our docking results, L105 is not essential for zafirlukast inhibition.

**Table 2.**
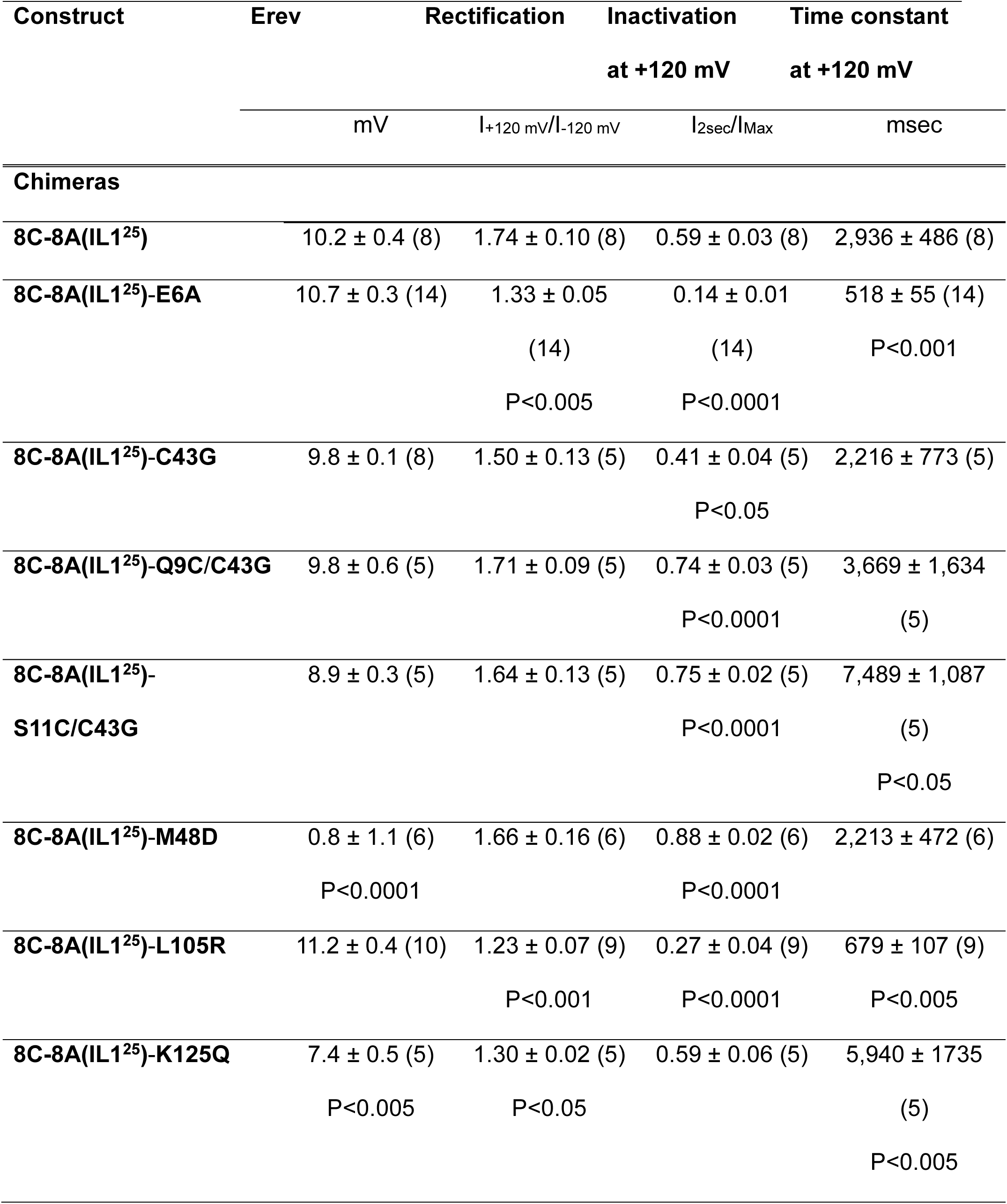
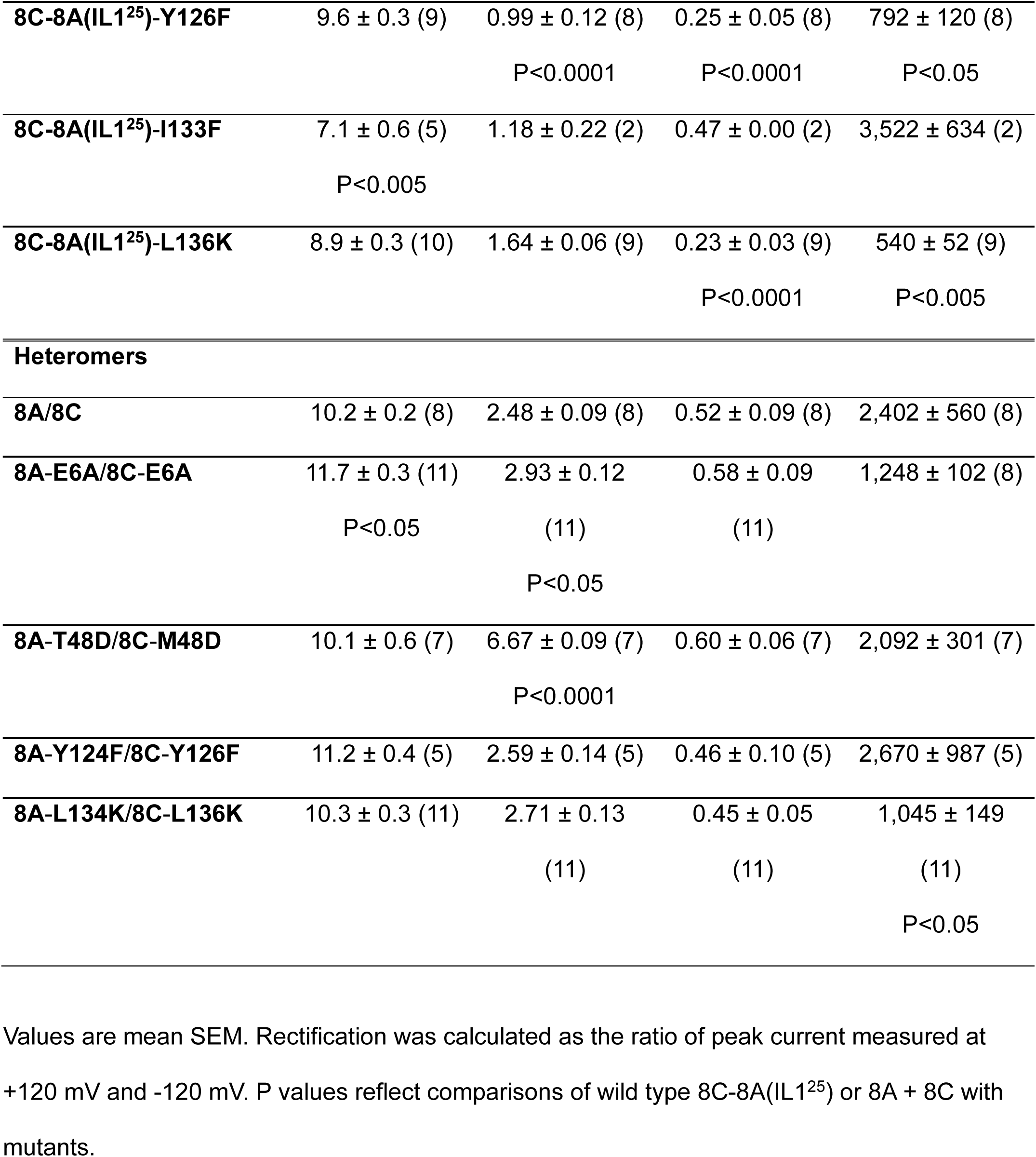
Electrophysiological properties of LRRC8 constructs.

| Construct | Erev | Rectification | Inactivation | Time constant |
| --- | --- | --- | --- | --- |
|  |  |  | at +120 mV | at +120 mV |
| | mV | $I_{+120 \text{ mV}}/I_{-120 \text{ mV}}$ | $I_{2\text{sec}}/I_{\text{Max}}$ | msec |
| **Chimeras** |  |  |  |  |
| **8C-8A(IL1<sup>25</sup>)** | 10.2 ± 0.4 (8) | 1.74 ± 0.10 (8) | 0.59 ± 0.03 (8) | 2,936 ± 486 (8) |
| **8C-8A(IL1<sup>25</sup>)-E6A** | 10.7 ± 0.3 (14) | 1.33 ± 0.05<br>(14)<br>P<0.005 | 0.14 ± 0.01<br>(14)<br>P<0.0001 | 518 ± 55 (14)<br>P<0.001 |
| **8C-8A(IL1<sup>25</sup>)-C43G** | 9.8 ± 0.1 (8) | 1.50 ± 0.13 (5) | 0.41 ± 0.04 (5)<br>P<0.05 | 2,216 ± 773 (5) |
| **8C-8A(IL1<sup>25</sup>)-Q9C/C43G** | 9.8 ± 0.6 (5) | 1.71 ± 0.09 (5) | 0.74 ± 0.03 (5)<br>P<0.0001 | 3,669 ± 1,634<br>(5) |
| **8C-8A(IL1<sup>25</sup>)-S11C/C43G** | 8.9 ± 0.3 (5) | 1.64 ± 0.13 (5) | 0.75 ± 0.02 (5)<br>P<0.0001 | 7,489 ± 1,087<br>(5)<br>P<0.05 |
| **8C-8A(IL1<sup>25</sup>)-M48D** | 0.8 ± 1.1 (6)<br>P<0.0001 | 1.66 ± 0.16 (6) | 0.88 ± 0.02 (6)<br>P<0.0001 | 2,213 ± 472 (6) |
| **8C-8A(IL1<sup>25</sup>)-L105R** | 11.2 ± 0.4 (10) | 1.23 ± 0.07 (9)<br>P<0.001 | 0.27 ± 0.04 (9)<br>P<0.0001 | 679 ± 107 (9)<br>P<0.005 |
| **8C-8A(IL1<sup>25</sup>)-K125Q** | 7.4 ± 0.5 (5)<br>P<0.005 | 1.30 ± 0.02 (5)<br>P<0.05 | 0.59 ± 0.06 (5) | 5,940 ± 1735<br>(5)<br>P<0.005 |
| **8C-8A(IL1<sup>25</sup>)-Y126F** | 9.6 ± 0.3 (9) | 0.99 ± 0.12 (8) | 0.25 ± 0.05 (8) | 792 ± 120 (8) |
|  |  | P<0.0001 | P<0.0001 | P<0.05 |
| **8C-8A(IL1<sup>25</sup>)-I133F** | 7.1 ± 0.6 (5) | 1.18 ± 0.22 (2) | 0.47 ± 0.00 (2) | 3,522 ± 634 (2) |
|  | P<0.005 |  |  |  |
| **8C-8A(IL1<sup>25</sup>)-L136K** | 8.9 ± 0.3 (10) | 1.64 ± 0.06 (9) | 0.23 ± 0.03 (9) | 540 ± 52 (9) |
|  |  |  | P<0.0001 | P<0.005 |
| **Heteromers** |  |  |  |  |
| **8A/8C** | 10.2 ± 0.2 (8) | 2.48 ± 0.09 (8) | 0.52 ± 0.09 (8) | 2,402 ± 560 (8) |
| **8A-E6A/8C-E6A** | 11.7 ± 0.3 (11) | 2.93 ± 0.12 | 0.58 ± 0.09 | 1,248 ± 102 (8) |
|  | P<0.05 | (11) | (11) |  |
|  |  | P<0.05 |  |  |
| **8A-T48D/8C-M48D** | 10.1 ± 0.6 (7) | 6.67 ± 0.09 (7) | 0.60 ± 0.06 (7) | 2,092 ± 301 (7) |
|  |  | P<0.0001 |  |  |
| **8A-Y124F/8C-Y126F** | 11.2 ± 0.4 (5) | 2.59 ± 0.14 (5) | 0.46 ± 0.10 (5) | 2,670 ± 987 (5) |
| **8A-L134K/8C-L136K** | 10.3 ± 0.3 (11) | 2.71 ± 0.13 | 0.45 ± 0.05 | 1,045 ± 149 |
|  |  | (11) | (11) | (11) |
|  |  |  |  | P<0.05 |

Values are mean SEM. Rectification was calculated as the ratio of peak current measured at +120 mV and -120 mV. P values reflect comparisons of wild type 8C-8A(IL1<sup>25</sup>) or 8A + 8C with mutants.

We next evaluated the zafirlukast sensitivity of 8C-8A(IL1^25^) chimeras, which carry mutations in the zafirlukast docking site formed by TM1 and TM2. Mutation in TM1 of cysteine 43 to glycine (C43G), the corresponding residue in 8A, 8B, and 8D, enhanced VDI (Fig. 4h and Table 2) and led to a ∼3-fold reduction in zafirlukast IC_50_ (Fig. 4o and Table 1). Methionine 48 was mutated to aspartate (M48D) because the analogous mutation in 8A-T48D/8C heteromers causes constitutive opening due to electrostatic repulsion of pore-blocking lipids^31^. Similarly, we found that 8C-8A(IL1^25^)-M48D is constitutively open and insensitive to cell swelling or shrinkage (Supplemental Fig. 1). VDI was largely absent in this mutant (Fig. 4i and Table 2). Despite being constitutively open, the M48D mutant is still sensitive to inhibition by zafirlukast, albeit with twofold lower sensitivity (Fig. 4o and Table 1).

The residues on TM2 comprising the zafirlukast docking site are likely surrounded by membrane phospholipids^26^, suggesting that channel-lipid interactions could impact zafirlukast pharmacology. Therefore, mutations were designed to disrupt zafirlukast binding as well as destabilize putative 8C- 8A(IL1^25^) channel-lipid interactions. These mutations are lysine 125-to-glutamine (K125Q), tyrosine 126-to-phenylalanine (Y126F), isoleucine 133-to-phenylalanine (I133F), and leucine 136-to-lysine (L136K).

K125Q (Fig. 4j and Table 2) and Y126F (Fig. 4k and Table 2) mutations significantly reduced the rectification properties of 8C-8A(IL1^25^) whereas Y126F (Fig. 4k and Table 2) and L136K (Fig. 4m and Table 2) mutants exhibited greater VDI. All four mutations significantly enhanced zafirlukast sensitivity (Fig. 4p and Table 1), with the L136K mutation reducing IC_50_ by approximately 90-fold.

### Role of the N-terminus in zafirlukast mechanism of action

The NTD was not resolved in 8C-8A(IL1^25^) cryo-EM structure^26^ or most published LRRC8 structures^27, 29, 33, 41–43^ due to its inherent structural flexibility. Two exceptions are 8A and 8D homomeric structures, which show that the NTD extends into the pore and creates a second constriction site ^28, 30^. When predicted using AlphaFold3^37^, the structural model of the 8C-8A(IL1^25^) chimera suggests that the NTD extends into the inner pore as observed in the structures of 8A and 8D and partially overlaps with the putative zafirlukast binding site (Fig. 5a-b). This model was tested in patch clamp experiments by evaluating if cysteines introduced into the NTD are able to coordinate cadmium (Cd^2+^) ions applied extracellularly to open channels. Optimal Cd^2+^ coordination by cysteines occurs when the residues are approximately ∼5Å apart. Five N-terminal residues (E6, R8, Q9, S11, and E12) were individually mutated to cysteine either alone or together with C43G, the latter of which removes an endogenous cysteine residue present in the pore of 8C. Single cysteine mutants and double R8C/C43G and E12C/C43G mutants generated either non-functional channels or small-amplitude currents under standard transfection conditions (data not shown). Q9C/C43G and S11C/C43G channels generated robust swelling-activated currents, however, cells transfected with 8C-8A(IL1^25^)-S11C/C43G were unhealthy and difficult to patch clamp (data not shown). Inspection of the 8C-8A(IL1^25^) structure identified an additional endogenous cysteine residue at position 141 (C141) that could potentially coordinate Cd^2+^, leading us to mutate this residue to serine (C141S) in the 8C-8A(IL1^25^)-C43G and 8C-8A(IL1^25^)-Q9C/C43G mutants. Both mutants were functional and amenable to experimentation. Following steady-state activation with cell swelling, bath addition of 100 µM Cd^2+^ had no effect on C43G/C141S mutants, whereas Cd^2+^ inhibited Q9C/C43G/C141S-mediated currents by approximately 50% (Fig. 5c). These data support our AlphaFold3 model suggesting the NTD extends into pore of 8C-8A(IL1^25^) and approaches the zafirlukast docking site.

**Figure 5.**
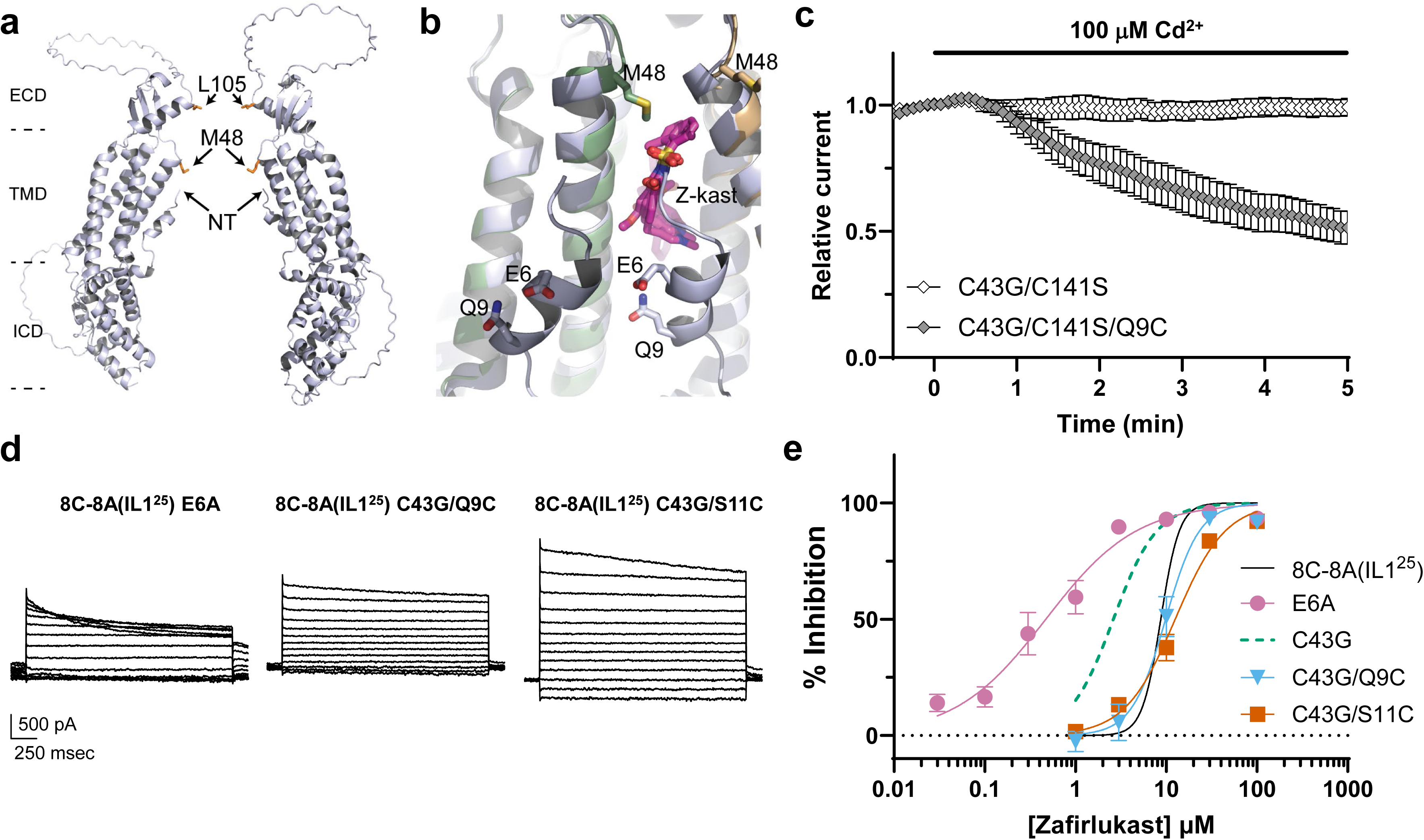
Role of the NTD in 8C-8A(IL1^25^) sensitivity to zafirlukast. AlphaFold3 model of the entire pore region of 8C-8A(IL1^25^), including the NTD overlayed on individual subunits of the cryo-EM structure of heptameric 8C-8A(IL1^25^) (PDB ID: 8DXN) to visualize the positioning of the NTD, which is not resolved in the cryo-EM structures. **a)** Opposing subunits are shown in ribbon representation to highlight the narrowing of the pore by the N-terminal residues. **b)** Close-up view of the putative zafirlukast binding site with the docked zafirlukast molecules. The AlphaFold model (light blue) is overlayed onto individual subunits (gold and green) that form the binding site. **c)** Coordination of Cd^2+^ by Q9C in the NTD. Cells transfected with C43G/C141S (control) or C43G/C141S/Q9C were subjected to hypotonic cell swelling and then treated with 100 µM Cd^2+^ applied to the bath. Data are mean ± SEM normalized current amplitude recorded at +100 mV (n=5 in each condition). **d)** Representative currents recorded from the indicated NTD mutants. **e)** Mean ± SEM CRC data for the indicated mutants (WT n = 4-8, E6A n = 4-7, C43G n = 5-6, C43G/Q9C n = 5, C43G/S11C n = 5). Fit of WT 8C-8A(IL1^25^) CRC data (black line) from Fig. 2c is shown for comparison.

We tested this model by evaluating the effects of NTD mutations on zafirlukast sensitivity. As noted earlier, the single C43G mutation enhanced VDI and zafirlukast sensitivity compared to WT 8C-8A(IL1^25^) (Fig. 4o and Table 1). Interestingly, the double Q9C/C43G and S11C/C43G mutations reversed the effects of the C43G mutation on both VDI and zafirlukast IC_50_ (Fig. 5d-e and Table 1). The E6A mutation increased both VDI and sensitivity to zafirlukast compared to WT 8C-8A(IL1^25^) (Fig. 5d-e and Table 1).

### The mechanism of zafirlukast inhibition is conserved in 8A/8C channels

To determine if the mechanism of zafirlukast action in 8C-8A(IL1^25^) homomeric channels is conserved in heteromers, we assessed the effects of select NTD, TM1, and TM2 mutations in 8A/8C. These were 8A-E6A/8C-E6A, 8A-T48D/8C-M48D, 8A-Y124F/8C-Y126F, and 8A-L134K/8C-L136K.

Zafirlukast inhibited WT 8A/8C currents with a significantly (P<0.0001) slower time constant (tau=400 sec at +100 mV) than that of 8C-8A(IL1^25^) (Fig. 6f) and IC_50_ values of 17.6 µM and 15.2 µM at +100 mV and -100 mV, respectively (Fig. 6g and Table 1). Unlike the effects of mutations in 8C-8A(IL1^25^), none of the 8A/8C mutations altered VDI (Fig. 6a-e and Table 2). However, consistent with the pattern observed with the chimera, E6A/E6A and L134K/L136K mutants were more sensitive to zafirlukast, whereas T48D/M48D channels exhibited a blunted response to the drug. The Y124F/Y126F mutations had no effect on zafirlukast sensitivity (Fig. 6g and Table 1). Taken together, these data indicate that the mechanism of zafirlukast action is largely conserved in LRRC8A/LRRC8C heteromeric channels.

**Figure 6.**
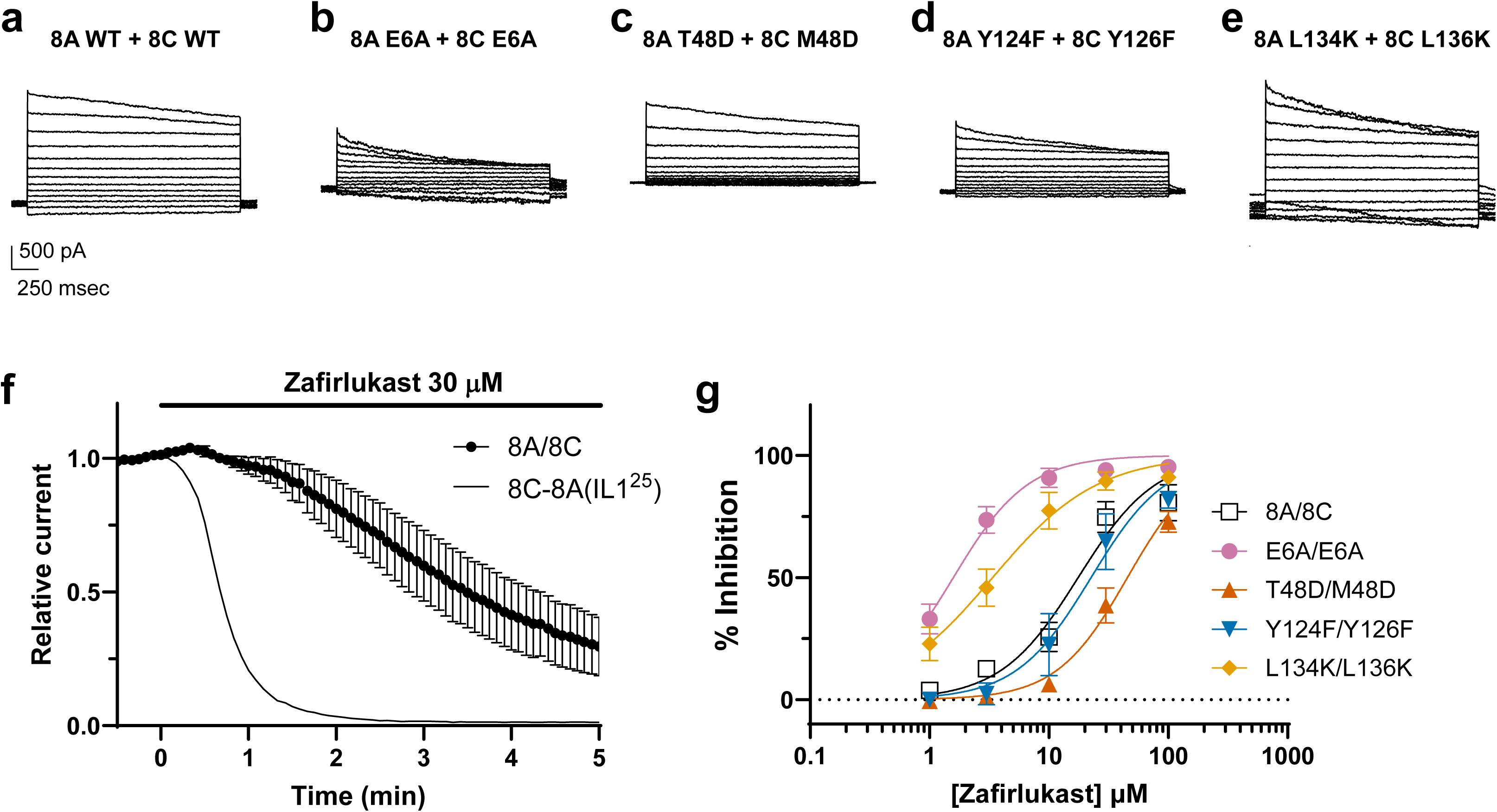
Analysis of docking site mutations on heteromeric LRRC8A/LRRC8C sensitivity to zafirlukast. **a-e)** Representative WT and mutant LRRC8A/LRRC8C currents. **f)** Time-course of LRRC8A/LRRC8C current inhibition by 30 µM zafirlukast. Data are means ± SEM (n=5). Time-course of 8C-8A(IL1^25^) inhibition by 30 µM from Fig. 2b is shown for comparison. **g)** Mean ± SEM CRC data for WT and mutant LRRC8A/LRRC8C at +100 mV (8A/8C n = 6-8, E6A/E6A n = 6, T48D/M48D n = 4-7, Y124F/Y126F n = 5, L134K/L136K n = 5).

### The mechanism of pranlukast inhibition is conserved in 8C-8A(IL1^25^) and 8A/8C channels

We previously reported that pranlukast, another CysLT1 receptor antagonist, also inhibits native VRACs independently of the receptor^22^. This was surprising given that the compounds are structurally distinct and in fact represent different chemical scaffolds (Fig. 1). One chemical-physical property they do share is a high partition coefficient, logP, owing to their high lipophilicity. Considering that zafirlukast interacts with a hydrophobic site in LRRC8 channels, we tested if pranlukast sensitivity is also altered by zafirlukast docking site mutations. Pranlukast inhibited WT 8C-8A(IL1^25^) with IC_50_ values of 1.5 µM and 5.9 µM at +100 mV and -100 mV, respectively. Single E6A, M48D, Y126F, and L136K mutations all significantly affected pranlukast sensitivity with the same directionality observed for zafirlukast (Fig. 7a and Table 3). Pranlukast inhibited heteromeric 8A/8C channels with IC_50_ values of 9.3 µM and 11.8 µM at +100 mV and -100 mV, respectively. Double E6A/E6A and L134K/L36K mutations significantly enhanced zafirlukast sensitivity, whereas Y124F/Y126F mutation did not. T48D/M48D channels were virtually insensitive to pranlukast (Fig. 7b and Table 3). The overall correspondence between zafirlukast and pranlukast sensitivity to these mutations suggests the drugs inhibit LRRC8 channels through a conserved mechanism of action.

**Figure 7.**
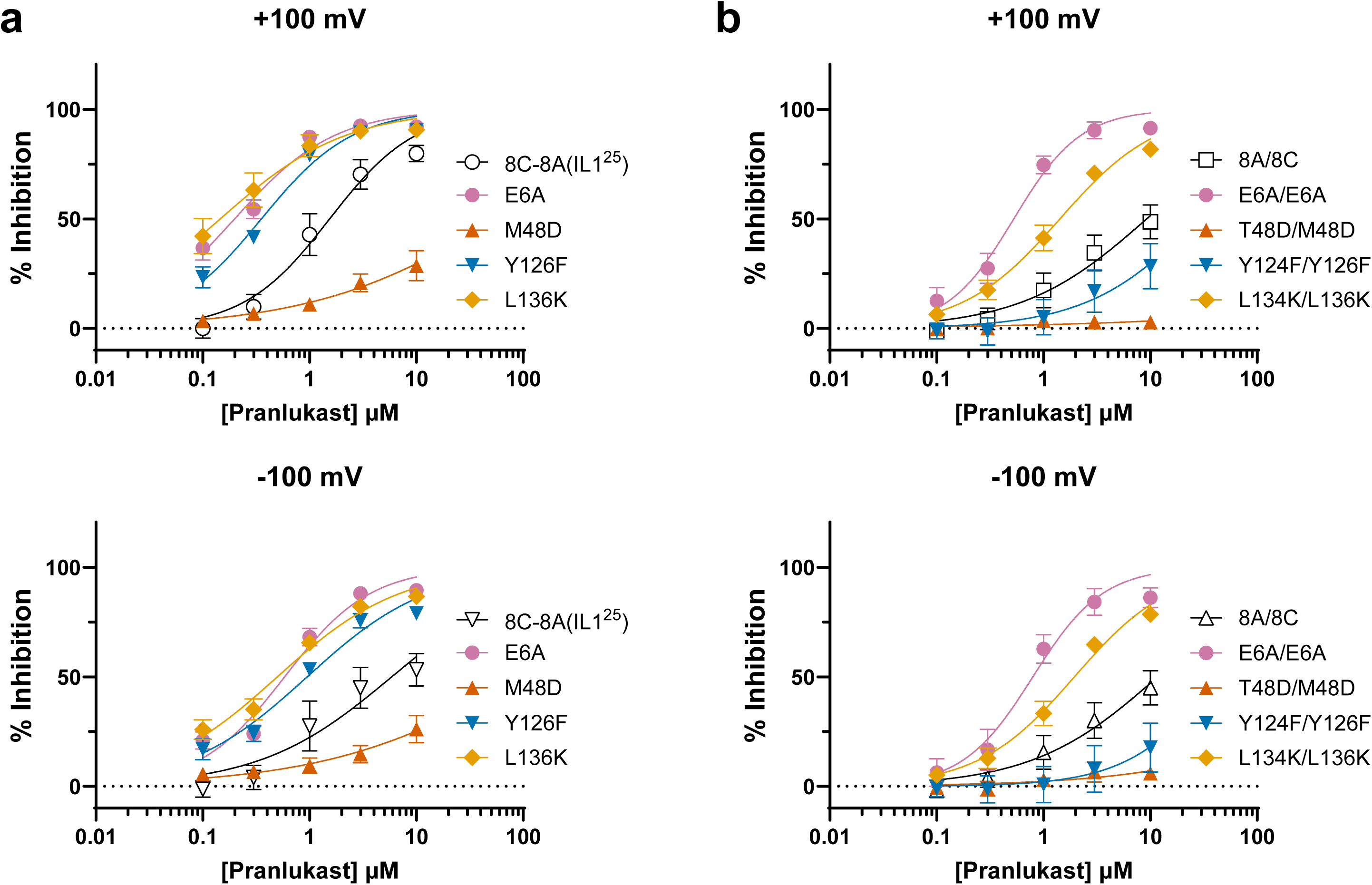
Analysis of docking site mutations on 8C-8A(IL1^25^) and heteromeric LRRC8A/LRRC8C sensitivity to pranlukast. **a-b)** Pranlukast CRC data for WT 8C-8A(IL1^25^) and the indicated mutants at +100 mV (top panel) and -100 mV (bottom panel). **c-d)** Pranlukast CRC data for WT LRRC8A/LRRC8C and the indicated mutants at +100 mV (top panel) and -100 mV (bottom panel). Data are means ± SEM (WT n = 4, E6A, n = 5, M48D n = 5, Y126F n = 5, L136K n = 5, 8A/8C n = 6, E6A/E6A n = 6, T48D/M48D n = 3, Y124F/Y126F n = 6, L134K/L136K n = 6).

**Table 3.** Pranlukast sensitivity of LRRC8 constructs.

| Construct | IC<sub>50</sub> (μM) |  | Hill coefficient |  |
| --- | --- | --- | --- | --- |
|  | +100 mV | -100 mV | +100 mV | -100 mV |
| **Chimera** |  |  |  |  |
| **8C-8A(IL1<sup>25</sup>)** | 1.5 | 5.9 | 1.08 | 0.70 |
| **8C-8A(IL1<sup>25</sup>)-E6A** | 0.20 | 0.58 | 0.93 | 1.08 |
|  | P<0.0001 | P<0.0001 |  |  |
| **8C-8A(IL1<sup>25</sup>)-M48D** | 57.6 | 100.1 | 0.50 | 0.47 |
|  | P<0.0001 | P<0.001 | P<0.05 |  |
| **8C-8A(IL1<sup>25</sup>)-Y126F** | 0.35 | 0.95 | 1.02 | 0.76 |
|  | P<0.0001 | P<0.0001 |  |  |
| **8C-8A(IL1<sup>25</sup>)-L136K** | 0.15 | 0.5 | 0.74 | 0.75 |
|  | P<0.0001 | P<0.0001 |  |  |
| **Heteromer** |  |  |  |  |
| **8A/8C** | 9.3 | 11.8 | 0.74 | 0.74 |
| **8A-E6A/8C-E6A** | 0.53 | 0.79 | 1.39 | 1.33 |
|  | P<0.0001 | P<0.0001 | P<0.05 |  |
| **8A-T48D/8C-M48D** | No effect | No effect | - | - |
| **8A-Y124F/8C-Y126F** | 28.6 | 45.1 | 0.82 | 1.00 |
| **8A-L134K/8C-L136K** | 1.4 | 1.9 | 0.95 | 0.96 |
|  | P<0.0001 | P<0.0001 |  |  |

P values reflect comparisons of wild type 8C-8A(IL1<sup>25</sup>) or 8A + 8C with mutants.

### Zafirlukast inhibition is not dependent on the lipid gate

As noted earlier, emerging structural and functional evidence indicate that membrane phospholipids play a direct role in LRRC8 channel gating in response to cell volume changes^26, 31^, raising the possibility that zafirlukast inhibits channel activity by closing the lipid gate. We therefore developed a Substituted Cysteine Accessibility Mutagenesis (SCAM) assay of functional MTSET reactivity for measuring lipid gate closure. MTSET was chosen as the thiol reactive reagent because its positive charge renders it membrane impermeant and restricted to aqueous solvent. We hypothesized that in the shrinkage-induced inactivated state, the channel pore would be occluded by phospholipids and impermeant to extracellular MTSET, whereas opening of the lipid gate during swelling would allow MTSET to traverse the aqueous pore and react with Q9C in the NTD (Fig. 8a). WT and C43G channels were inhibited by approximately 40% following bath application of 1 mM MTSET for 3 min (Supplemental Fig. 2), whereas Q9C/C43G mutants were inhibited by approximately 80%, which is significantly (P<0.05) greater than that observed for either WT or C43G channels. We therefore used Q9C/C43G mutants for SCAM experiments. Q9C/C43G channels were first maximally closed by hypertonic shrinkage, subsequently activated with hypotonic swelling, and then treated with MTSET. Under these conditions, MTSET almost fully inhibited swelling-activated Q9C/C43G currents (Fig. 8b). Importantly, however, pre-application of MTSET to inactivated channels had no significant effect on the magnitude of MTSET-inhibitable swelling-activated Q9C/C43G current (Fig. 8c-d). These data suggest MTSET reactivity with Q9C/C43G can be used as a probe of the lipid gate.

**Figure 8.**
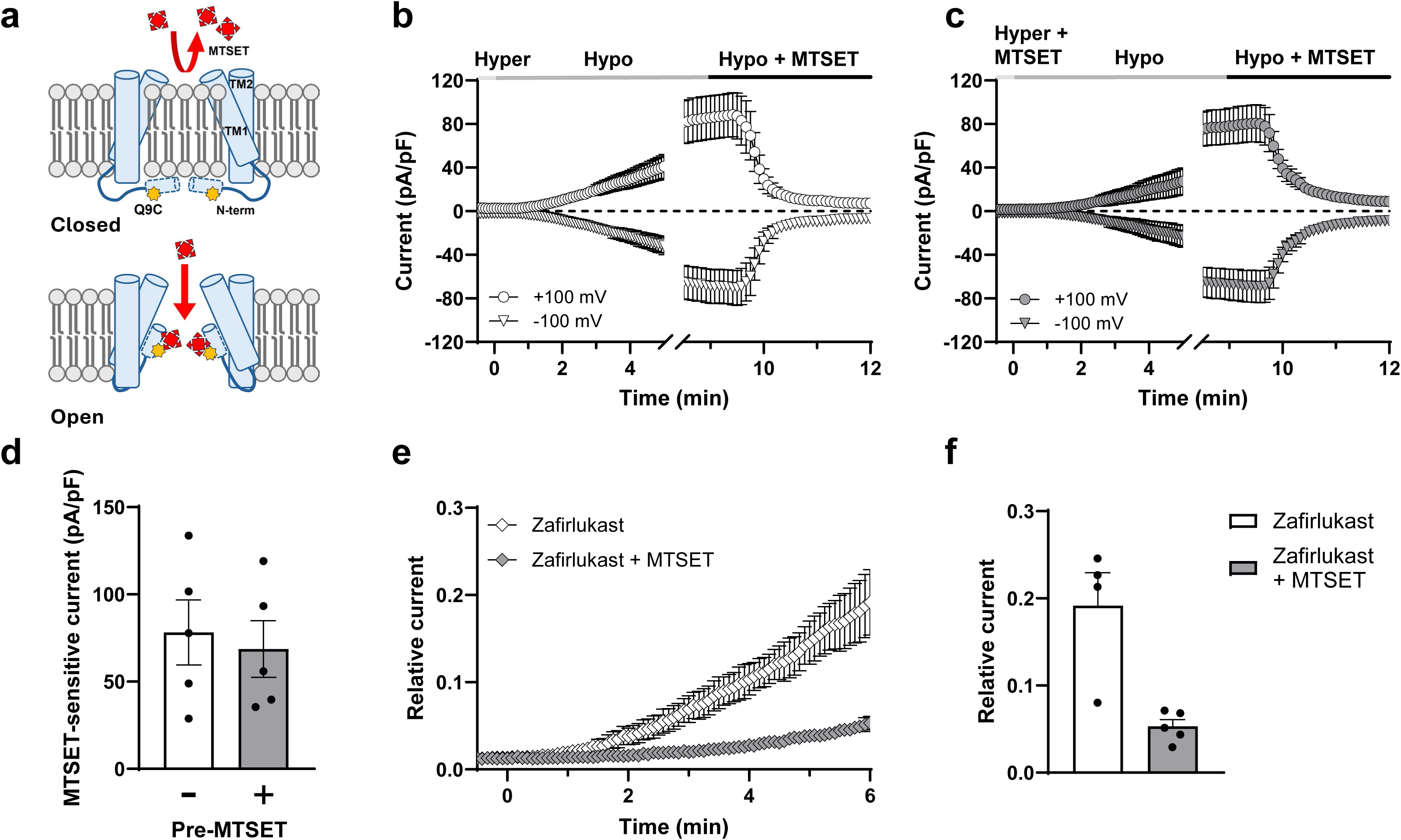
Zafirlukast does not close the lipid gate. **a)** hypothetical models of MTSET reaction. **b)** Inhibition of swelling-activated 8C-8A(IL1^25^)-Q9C/C43G currents with MTSET. Cells were treated with hypertonic saline for 15 min to fully close 8C-8A(IL1^25^)-Q9C/C43G channels, subjected to hypotonic cell swelling for 9 min to maximally activate 8C-8A(IL1^25^)-Q9C/C43G currents, and then treated with 1 mM MTSET in hypotonic buffer for 3 min to inhibit 8C-8A(IL1^25^)-Q9C/C43G. Data are mean ± SEM current density at +100 mV (open circles) and -100 mV (open inverted triangles; n=5). **c)** Closure of the lipid gate protects 8C-8A(IL1^25^)-Q9C/C43G currents from MTSET reactivity. Cells were treated with hypertonic saline for 8 min, treated for 3 min with hypertonic saline containing 1 mM MTSET, washed for 4 min with hypertonic buffer, subjected to hypotonic cell swelling for 9 min to maximally activate 8C-8A(IL1^25^)-Q9C/C43G currents, and then treated with 1 mM MTSET in hypotonic buffer for 3 min to inhibit 8C-8A(IL1^25^)-Q9C/C43G. Data are mean ± SEM current density at +100 mV (closed circles) and -100 mV (closed inverted triangles; n=5). **d)** Mean ± SEM MTSET-inhibitable current density at +100 mV in cells pre-treated with hypertonic buffer alone (open bars) or with hypertonic buffer containing 1 mM MTSET (grey bars; n=5). **e)** Zafirlukast inhibition does not protect 8C-8A(IL1^25^)-Q9C/C43G currents from MTSET reactivity. Cells were dialyzed with low intracellular ionic strength pipette solution to activate 8C-8A(IL1^25^)-Q9C/C43G currents, pre-treated with 30 µM zafirlukast alone (3 min) to fully inhibit channel activity, switched to buffer containing either zafirlukast alone (open symbols) or together with 1 mM MTSET (closed symbols) for 3.5 min, and then washed with control isotonic buffer. Data are mean ± SEM current density recorded at +100 mV (n=4-5). **f)** Fraction of recovered current density after 6 min of washout (n=4-5). *P<0.005.

We next asked if Q9C/C43G inhibition by zafirlukast involves closure of the lipid gate. We took advantage of the partial washout of zafirlukast following channel activation with low intracellular ionic strength (Fig. 2e) and determined if zafirlukast inhibition protects Q9C/C43G from MTSET reactivity. Similar to WT 8C-8A(IL1^25^), zafirlukast inhibition was partially reversible in Q9C/C43G channels activated with low ionic strength. However, no current recovery was observed when MTSET was co-applied during zafirlukast inhibition (Fig. 8e-f). Taken together, these data suggest that whereas shrinkage-induced channel inactivation involves closure of the lipid gate, the zafirlukast inhibition does not.

### Zafirlukast inhibits LRRC8/VRAC by promoting channel inactivation

We previously reported that pranlukast enhances VDI in native VRAC currents^22^, prompting us to examine the effects of zafirlukast on 8C-8A(IL1^25^) VDI here. The time course of inactivation at +120 mV was well fit by a single exponential. As shown in Fig. 9a-b, zafirlukast, administered at an IC_50_ dose of 9 µM, significantly (P<0.05) reduced the tau of VDI by approximately 50%. Zafirlukast did not affect 8C-8A(IL1^25^) reactivation at -120 mV (data not shown). We noted that many of the NTD, TM1, and TM2 mutants exhibiting greater sensitivity to zafirlukast also showed greater VDI. Plotting mean zafirlukast IC_50_ against VDI (I_2sec_/I_max_) revealed a significant correlation, suggesting a mechanistic link between the two. We explored this possibility by testing if enhancement of VDI in native VRAC currents with low extracellular pH^40, 44^ also increases zafirlukast sensitivity. Wild type HCT cells expressing all 5 *LRRC8* genes^5^ were voltage clamped between +40 and +120 mV to induce VDI. As shown in Fig. 9d-e, VDI is dramatically increased by switching extracellular pH from 7.4 to pH 6.0. Additionally, the zafirlukast CRC was significantly left-shifted in low pH conditions (IC_50_ pH 7.4 = 13.5 μM; IC_50_ pH 6.0 = 5.6 μM). Taken together, these data support a model in which zafirlukast inhibits LRRC8/VRAC by stabilizing an inactivated state of the channel.

**Figure 9.**
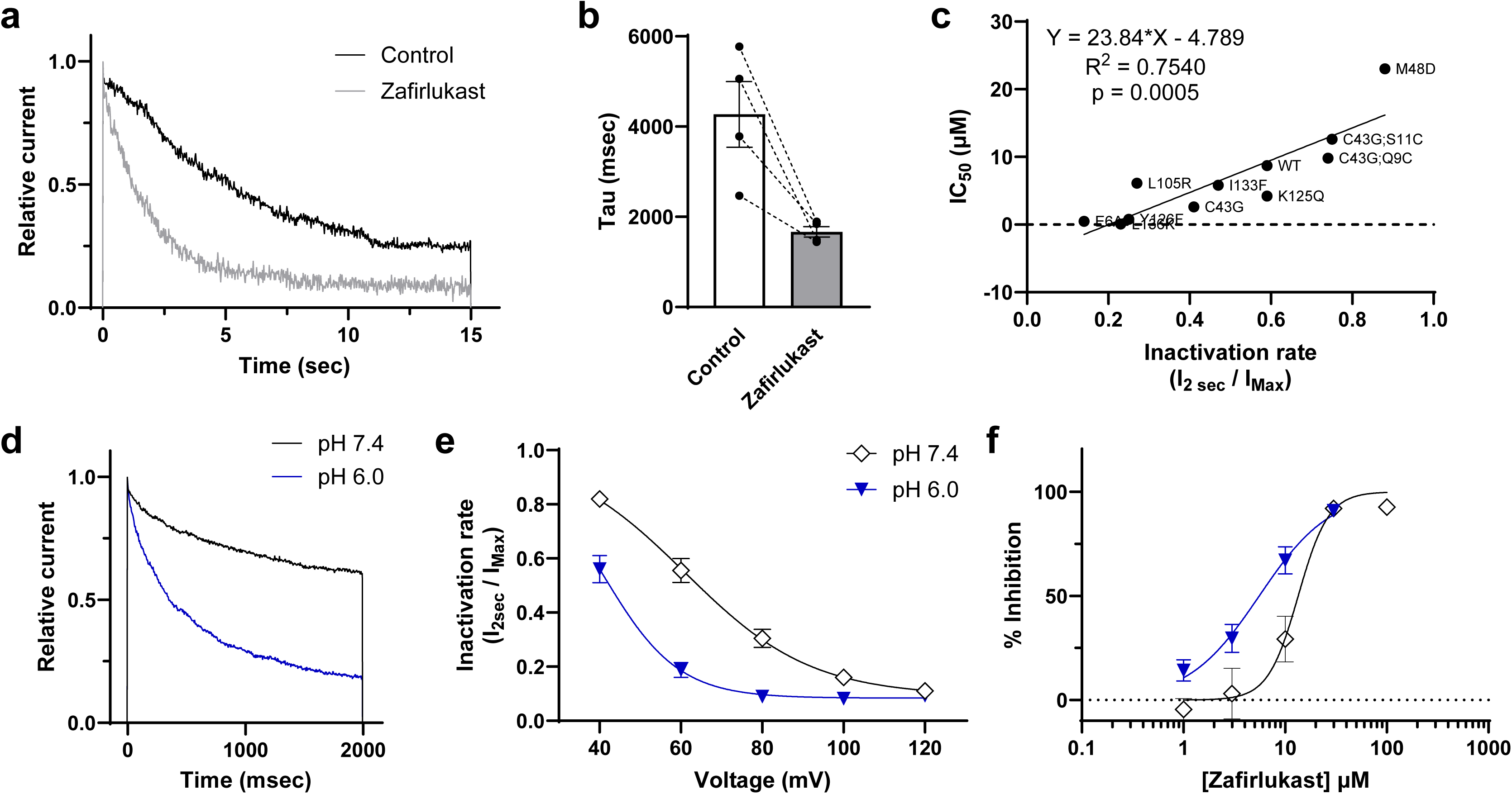
Relationship between VDI and zafirlukast sensitivity. **a)** 8C-8A(IL1^25^) currents were evoked at +120 mV to induce VDI in the absence (black trace) and presence (grey trace) of 9 µM zafirlukast. **b)** Time constants of VDI derived from single exponential fits. **c)** Plot of mean inactivation rate (I_2sec_/I_max_) vs IC_50_ for the indicated mutants. Linear fit data are shown in the inset. **d)** Representative traces recorded at +60 mV from wild type HCT cells at extracellular pH 7.4 (black trace) and pH 6.0 (blue trace). **e)** pH-induced enhancement of VDI. **f)** Zafirlukast CRC data recorded at +100 mV from HCT cells at pH 7.4 (open symbols) and pH 6.0 (blue symbols). Data are means ± SEM (n=5).

## DISCUSSION

Here we describe a conserved mechanism of LRRC8/VRAC inhibition by two structurally distinct CysLT1R antagonists that inhibit the channel independently of receptor signaling ^22^. In silico docking and AlphaFold modeling identified a putative zafirlukast binding site in 8C-8A(IL1^25^) sandwiched between the NTD, TM1, and TM2 near inter-subunit fenestrae believed to interact closely with plasma membrane phospholipids. In support of these models, mutations in the zafirlukast site led to variable and, in some cases, striking changes in zafirlukast and pranlukast pharmacology of both 8C-8A(IL1^25^) and 8A/8C channels. The proportional relationship between VDI and inhibitor sensitivity supports the idea that both drugs promote channel inactivation.

The zafirlukast docking site between subunits theoretically allows for a maximum of seven drug molecules to bind to each heptamer. The observed Hill coefficients > 1.0 for WT 8C-8A(IL1^25^) and 8A/8C are consistent with multiple ligands binding in a cooperative manner during channel inhibition. The molecular basis for this cooperativity is currently unknown but may be related to drug-induced destabilization of lipid-channel interactions (see below).

Zafirlukast docks near C43 of TM1 and K125, Y126, I133, and L136 of TM2. With the exception of M48, none of the side chains projects into conduction pore and are therefore unlikely to directly contribute to anion binding and conduction. On the contrary, we believe these residues play important roles in phospholipid interactions with the channel. M48 likely stabilizes lipids that occlude the pore in the closed state, as the introduction of negative charge at this site (i.e. 8C-8A(IL1^25^)-M48D and 8A-T48D/8C-M48D) leads to constitutive channel activation (this study and ref.^31^). None of the other TM1 or TM2 mutants were constitutively active (data not shown). K125 and Y126 side chains project into the fenestrae and may interact with surrounding polar lipid head groups. I133 and L136 also extend into the fenestrae and may stabilize TM2 interactions with lipid acyl chains.

We were surprised that every mutation we tested in 8C-8A(IL1^25^) apart from M48D led to an increase in drug sensitivity, suggesting a common underlying mechanism. There are at least 3 possible explanations, which are not mutually exclusive. First, it is conceivable, albeit unlikely in our view, that all of the sensitizing mutations increase the binding site affinity for both structurally distinct drugs. Zafirlukast and pranlukast are single-nanomolar-affinity orthostatic antagonists of CysLT1R, which belongs to the heptahelical transmembrane domain family of lipid G-protein-coupled receptors^24^. They terminate signaling by displacing the endogenous inflammatory ligand, leukotriene D4 (LTD4), from its binding pocket^23^. Despite being chemically and structurally very distinct, both drugs occupy the same general binding pocket through multiple conserved and non-conserved contact points comprising hydrophobic, polar, hydrogen bond, and salt bridge interactions. One conserved interaction involves CysLT1R-Y104, which makes polar interactions with the tetrazol and benzopyran groups of pranlukast and sulfonamide moiety of zafirlukast^24^. Evaluating the structure-activity relationships (SAR) of zafirlukast and pranlukast analogs containing substitutions of these and other functional groups will be helpful in identifying the chemical and physical properties underlying channel inhibition. These studies may help inform the design of more potent and selective LRRC8/VRAC inhibitors.

Second, the zafirlukast docking site in LRRC8 is likely occupied by membrane phospholipids in the absence of the drug, raising the possibility that the inhibitors must first displace lipids from the channel TM domains before they can bind. This is analogous to how the antagonists displace LTD4 from CysLT1R^45^. Mutagenesis analysis has revealed an inverse relationship between LTD4 and antagonist potencies, such that mutations that weaken LTD4 binding reciprocally enhance zafirlukast and pranlukast potency, and vice-versa^24^. Similarly, we found that mutations in LRRC8 predicted to weaken lipid interactions also enhanced drug sensitivity. The L136K mutation, for example, which is predicted to be highly disruptive of lipid interactions, caused the greatest enhancement of drug sensitivity, whereas more subtle mutations (e.g., K125Q and I133F), had smaller effects. Evaluating how different types and positions of mutations along TM1 and TM2 alter drug sensitivity should help clarify the relationship between lipid binding and drug sensitivity. It should be noted that the docking computations identifying the zafirlukast binding site were performed against the apo structure of 8C-8A(IL1^25^) without any associated phospholipids. It will be important to evaluate how the inclusion of lipids in the docking model alters zafirlukast binding.

Finally, lipid packing in fenestrae and around TMs may impart structural rigidity to the transmembrane pore of LRRC8 channels. Drug binding and drug-sensitizing mutations could destabilize lipid interactions leading to greater structural flexibility, pore instability, and channel inactivation. Several lines of evidence support this idea. First, both zafirlukast (this study) and pranlukast^22^ enhance VDI, which is postulated to result from constriction of the outer pore^40, 44, 46^. Second, drug-sensitizing mutations predicted to destabilize lipid interactions enhance VDI. Third, the M48D mutation in 8C-8A(IL1^25^) decreased both drug sensitivity and VDI. Given that analogous mutations in LRRC8A/LRRC8C produce constitutively open channels free of pore-occluding lipids, it is possible these mutations increase lipid packing in fenestrae, increase structural rigidity and open-state stability, and reduce drug-induced inactivation. Fourth, low pH conditions that enhance VDI^40, 44, 47^ also increase zafirlukast sensitivity. Kinetic modeling of low pH effects on native VRAC has suggested that protonation of extracellular residues leads to destabilization and collapse of pore^44^. It is interesting to note that exogenous application of the lipid molecule arachidonic acid enhances VDI as part of VRAC inhibition independently of cyclooxygenase signaling^48^. Collectively, these data support the idea that modulatory drugs and membrane lipids regulate VRAC through interactions with channel fenestrations. Cryo-EM analysis of drug-sensitizing mutants should help illuminate the relationships between channel-lipid interactions, VDI, and pharmacology.

The intracellular NTD of 8C-8A(IL1^25^) was not resolved in the cryo-EM structure^26^ owing to its structural flexibility. We demonstrated here with computational and functional experiments that the NTD extends deep into the transmembrane pore, similar to 8A^30^ and 8D^28^ homomeric channels. MTSET reactivity experiments suggest that Q9C in the NTD is located on the intracellular side of the lipid gate of closed channels and that cell swelling-induced activation is associated with de-lipidation of the pore. The ability of Q9C to coordinate Cd^2+^ suggests these residues are within 5Å of each other in the open state^49^. In the AlphaFold3 model, the NTD of 8C-8A(IL1^25^) appears to partially overlap and perhaps even obscure the zafirlukast docking site. Zafirlukast and pranlukast access their binding pocket in CysLT1R from the extracellular side of the receptor through either a small aqueous pore or a lateral entrance from the plasma membrane^24^. Similarly, we found that zafirlukast inhibits 8C-8A(IL1^25^) channels only when applied to the extracellular side of the channel. The narrow pore constriction created by the NTD (this study and refs.^30, 50^) and putative outer-facing pocket created by the NTD, TM1, and TM2 might explain the sidedness of zafirlukast action.

Zafirlukast inhibition of 8C-8A(IL1^25^) was partially reversible under low intracellular ionic strength conditions. Cytoplasmic LRR domains are involved in environmental sensing and undergo conformational motions associated with channel gating ^33, 51, 52^. Brohawn and colleagues noted in 8A/8C heteromers that the outward displacement of LRR domains is associated with the widening of fenestrae between subunits^31^. The intracellular NTD is also believed to undergo conformational changes in low ionic strength conditions^30^. Thus, conformational changes in the NTD and TMs surrounding the zafirlukast binding site may account for the drug washout observed with low intracellular ionic strength.

Although the NTD is an integral part of ion conduction, selectivity, and pore stability^50^, it is understudied due to the difficulties of capturing its flexible structure with cryo-EM and extreme sensitivity to mutations in functional studies. Recent molecular dynamics calculations identified an energy well favoring Cl^-^ binding to E6 in the NTD^30^, which is conserved across all LRRC8 subunits. Neutralizing the charge on E6 enhances VDI in 8C-8A(IL1^25^)-E6A and 8A-E6A/8C-E6A channels (this study and ref.^50^), consistent with a role in anion binding and stabilization of the pore. Furthermore, 8C-8A(IL1^25^)-E6A and 8A-E6A/8C-E6A channels are more sensitive to zafirlukast and pranlukast. E6 is predicted to face the aqueous pore, however, the hydrophobic E6A mutation may induce reorientation of the flexible NTD, destabilization of Cl^-^ binding and pore integrity, tendency toward inactivation, and consequently greater drug sensitivity. Zafirlukast inhibition does not appear to depend on the closure of the lipid gate because 1) the drug inhibits 8C-8A(IL1^25^)-M48D and 8A-T48/8C-M48D, and 2) inhibition does not protect Q9C from MTSET reactivity. However, it is possible that protection of Q9C from MTSET reactivity in closed channels is due to as-yet discovered mechanisms.

There were several notable differences between 8C-8A(IL1^25^) and 8A/8C pharmacology and sensitivity to mutations. For instance, 8C-8A(IL1^25^) is at least twice as sensitive to zafirlukast and pranlukast than 8A/8C. Because the only difference between 8C-8A(IL1^25^) and 8C is a 25-amino acid substitution from 8A in the 8C-IL domain, it is plausible that the inclusion of 8A modulates channel sensitivity to inhibitors. This raises the possibility of developing subtype-selective inhibitors that can distinguish between channels containing different ratios of 8A to other LRRC8 subunits^27^. It will be critical to investigate the drug sensitivity of 8A, 8C, 8D, and 8E chimeras^34^ as well as different heteromers for hints of subtype selectivity. We also found that unlike 8C-8A(IL1^25^), none of the sensitizing mutations enhanced VDI in 8A/8C. This may be because the 8A subunit is more rigid than 8C^27^ and resistant to conformation-dependent inactivation. The highly cooperative Hill coefficient of 8C-8A(IL1^25^) inhibition by zafirlukast (∼4) compared to that of 8A/8C (∼1) is consistent with greater 8C flexibility and susceptibility to drug-induced inactivation. A prediction of this model is that channel subtypes that are more susceptible to inactivation will also be more sensitive to drug inhibition, suggesting it might be possible to develop subtype-specific inhibitors that exploit a channel’s intrinsic pore stability.

Although zafirlukast and pranlukast represent entirely different chemical scaffolds, they have in common a high partition coefficient (LogP) due to their high lipophilicity. For reference, LogP values of most clinically used drugs range between 1.3 and 2.2, with higher values reflecting greater lipophilicity. The LogP of zafirlukast and pranlukast are 5.5 and 4.2, respectively. This raises important questions about the broader generalizability of the inhibitory mechanism identified here, as it applies to other inhibitors. The literature is replete with examples of how commonly used VRAC inhibitors that are all highly lipophilic, such as niflumic acid (LogP = 3.7), DIDS (LogP = 4.5), NPPB (LogP = 4.1), glibenclamide (LogP = 4.8), tamoxifen (LogP = 7.1), verapamil (LogP = 3.8), flufenamic acid (LogP = 5.2), and DCPIB (LogP = 6.8), enhance VDI during channel inhibition^,47, 53^. It will be important to determine if these drugs also inhibit LRRC8/VRAC channels through the gating mechanism identified here.

In conclusion, we have identified a conserved mechanism of LRRC8/VRAC channel inhibition by two chemically distinct drugs. The strong correspondence between drug sensitivity and inactivation suggests they are mechanistically linked. We propose that drug binding in lipophilic fenestrae leads to the weakening of lipid interactions with the channel, destabilization of the outer TMs, and constriction of the pore. The contributions of the NTD, TMs, and extracellular loop-helix domains to drug inhibition await further study. Zafirlukast and pranlukast should be useful tools for probing the molecular basis of drug-induced inactivation gating in VRACs. Structurally defined homomeric LRRC8 chimeras like 8C-8A(IL1^25^) will be valuable models for exploring VRAC structure-function relationships and molecule pharmacology.

## ACKNOWLEDGEMENTS

This work was supported by R01DK051610 to JSD.

## AUTHOR CONTRIBUTIONS

*Designed experiments:* TY, PB, EK and JSD

*Performed experiments and data analysis:* TY and PB

*Wrote, edited, and approved the final version of the manuscript:* TY, PB, EK and JSD

## REFERENCES

1. Strange, K., Yamada, T. & Denton, J.S. A 30-year journey from volume-regulated anion currents to molecular structure of the LRRC8 channel. J Gen Physiol 151, 100–117 (2019).

2. Nilius, B. & Droogmans, G. Ion channels and their functional role in vascular endothelium. Physiol Rev 81, 1415–1459 (2001).

3. Pedersen, S.F., Klausen, T.K. & Nilius, B. The identification of a volume-regulated anion channel: an amazing Odyssey. Acta Physiol (Oxf) 213, 868–881 (2015).

4. Okada, Y. Physiology of the volume-sensitive/regulatory anion channel VSOR/VRAC. Part 1: from its discovery and phenotype characterization to the molecular entity identification. J Physiol Sci **74**, 3 (2024).

5. Voss, F.K. et al. Identification of LRRC8 heteromers as an essential component of the volume-regulated anion channel VRAC. Science 344, 634–638 (2014).

6. Qiu, Z. et al. SWELL1, a plasma membrane protein, is an essential component of volume-regulated anion channel. Cell 157, 447–458 (2014).

7. Choi, H., Rohrbough, J.C., Nguyen, H.N., Dikalova, A. & Lamb, F.S. Oxidant-resistant LRRC8A/C anion channels support superoxide production by NADPH oxidase 1. J Physiol 599, 3013–3036 (2021).

8. Concepcion, A.R. et al. The volume-regulated anion channel LRRC8C suppresses T cell function by regulating cyclic dinucleotide transport and STING-p53 signaling. Nat Immunol 23, 287–302 (2022).

9. Zhou, C. et al. Transfer of cGAMP into Bystander Cells via LRRC8 Volume-Regulated Anion Channels Augments STING-Mediated Interferon Responses and Anti-viral Immunity. Immunity 52, 767–781 e766 (2020).

10. Wu, X. et al. The volume regulated anion channel VRAC regulates NLRP3 inflammasome by modulating itaconate efflux and mitochondria function. Pharmacol Res 198, 107016 (2023).

11. Wang, L. et al. ATP-elicited Cation Fluxes Promote Volume-regulated Anion Channel LRRC8/VRAC Transport cGAMP for Antitumor Immunity. J Immunol 213, 347–361 (2024).

12. Kang, C. et al. SWELL1 is a glucose sensor regulating beta-cell excitability and systemic glycaemia. Nat Commun 9, 367 (2018).

13. Gunasekar, S.K. et al. Small molecule SWELL1 complex induction improves glycemic control and nonalcoholic fatty liver disease in murine Type 2 diabetes. Nat Commun 13, 784 (2022).

14. Yang, J. et al. Glutamate-Releasing SWELL1 Channel in Astrocytes Modulates Synaptic Transmission and Promotes Brain Damage in Stroke. Neuron 102, 813–827 e816 (2019).

15. Chu, J. et al. ATP-releasing SWELL1 channel in spinal microglia contributes to neuropathic pain. Sci Adv 9, *eade9931* (2023).

16. Afzal, A. et al. The LRRC8 volume-regulated anion channel inhibitor, DCPIB, inhibits mitochondrial respiration independently of the channel. Physiol Rep 7, e14303 (2019).

17. Bowens, N.H., Dohare, P., Kuo, Y.H. & Mongin, A.A. DCPIB, the proposed selective blocker of volume-regulated anion channels, inhibits several glutamate transport pathways in glial cells. Mol Pharmacol 83, 22–32 (2013).

18. Deng, W., Mahajan, R., Baumgarten, C.M. & Logothetis, D.E. The ICl,swell inhibitor DCPIB blocks Kir channels that possess weak affinity for PIP2. Pflugers Arch 468, 817–824 (2016).

19. Lv, J. et al. DCPIB, an Inhibitor of Volume-Regulated Anion Channels, Distinctly Modulates K2P Channels. ACS Chem Neurosci **10**, 2786-2793 (2019).

20. Fujii, T. et al. Inhibition of gastric H+,K+-ATPase by 4-(2-butyl-6,7-dichloro-2-cyclopentylindan-1-on-5-yl)oxybutyric acid (DCPIB), an inhibitor of volume-regulated anion channel. Eur J Pharmacol **765**, 34-41 (2015).

21. Figueroa, E.E. & Denton, J.S. A SWELL time to develop the molecular pharmacology of the volume-regulated anion channel (VRAC). Channels (Austin) 16, 27–36 (2022).

22. Figueroa, E.E., Kramer, M., Strange, K. & Denton, J.S. CysLT1 receptor antagonists pranlukast and zafirlukast inhibit LRRC8-mediated volume regulated anion channels independently of the receptor. Am J Physiol Cell Physiol 317, C857–C866 (2019).

23. Tamada, T. & Ichinose, M. Leukotriene Receptor Antagonists and Antiallergy Drugs. Handb Exp Pharmacol 237, 153–169 (2017).

24. Luginina, A. et al. Structure-based mechanism of cysteinyl leukotriene receptor inhibition by antiasthmatic drugs. Sci Adv 5, eaax2518 (2019).

25. Yanushkevich, S. et al. Recent advances in the structure, function and regulation of the volume-regulated anion channels and their role in immunity. J Physiol (2024).

26. Takahashi, H., Yamada, T., Denton, J.S., Strange, K. & Karakas, E. Cryo-EM structures of an LRRC8 chimera with native functional properties reveal heptameric assembly. eLife 12 (2023).

27. Rutz, S., Deneka, D., Dittmann, A., Sawicka, M. & Dutzler, R. Structure of a volume-regulated heteromeric LRRC8A/C channel. Nat Struct Mol Biol 30, 52–61 (2023).

28. Nakamura, R. et al. Cryo-EM structure of the volume-regulated anion channel LRRC8D isoform identifies features important for substrate permeation. Commun Biol 3, 240 (2020).

29. Kefauver, J.M. et al. Structure of the human volume regulated anion channel. eLife 7 (2018).

30. Liu, H. et al. Structural insights into anion selectivity and activation mechanism of LRRC8 volume-regulated anion channels. Cell reports 42, 112926 (2023).

31. Kern, D.M. et al. Structural basis for assembly and lipid-mediated gating of LRRC8A:C volume-regulated anion channels. Nat Struct Mol Biol 30, 841–852 (2023).

32. Karakas, E., Strange, K. & Denton, J.S. Recent advances in structural characterization of volume-regulated anion channels (VRACs). J Physiol (2025).

33. Kern, D.M., Oh, S., Hite, R.K. & Brohawn, S.G. Cryo-EM structures of the DCPIB-inhibited volume-regulated anion channel LRRC8A in lipid nanodiscs. eLife 8 (2019).

34. Yamada, T. & Strange, K. Intracellular and extracellular loops of LRRC8 are essential for volume-regulated anion channel function. J Gen Physiol 150, 1003–1015 (2018).

35. Yamada, T., Figueroa, E.E., Denton, J.S. & Strange, K. LRRC8A homohexameric channels poorly recapitulate VRAC regulation and pharmacology. Am J Physiol Cell Physiol 320, C293–C303 (2021).

36. Chemical Computing Group, I. (Montreal, QC, Canada 2015).

37. Abramson, J. et al. Accurate structure prediction of biomolecular interactions with AlphaFold 3. Nature 630, 493–500 (2024).

38. McCann, J.D., Li, M. & Welsh, M.J. Identification and regulation of whole-cell chloride currents in airway epithelium. J Gen Physiol 94, 1015–1036 (1989).

39. Worrell, R.T., Butt, A.G., Cliff, W.H. & Frizzell, R.A. A volume-sensitive chloride conductance in human colonic cell line T84. Am J Physiol 256, C1111–1119 (1989).

40. Jackson, P.S. & Strange, K. Characterization of the voltage-dependent properties of a volume-sensitive anion conductance. J Gen Physiol 105, 661–676 (1995).

41. Kern, D.M. et al. Structural basis for assembly and lipid-mediated gating of LRRC8A:C volume-regulated anion channels. Nat Struct Mol Biol (2023).

42. Quinodoz, M. et al. De novo variants in LRRC8C resulting in constitutive channel activation cause a human multisystem disorder. EMBO J 44, 413–436 (2025).

43. Kasuya, G. et al. Cryo-EM structures of the human volume-regulated anion channel LRRC8. Nat Struct Mol Biol 25, 797–804 (2018).

44. Hernandez-Carballo, C.Y., De Santiago-Castillo, J.A., Rosales-Saavedra, T., Perez-Cornejo, P. & Arreola, J. Control of volume-sensitive chloride channel inactivation by the coupled action of intracellular chloride and extracellular protons. Pflugers Arch 460, 633–644 (2010).

45. Ravasi, S., Capra, V., Mezzetti, M., Nicosia, S. & Rovati, G.E. A kinetic binding study to evaluate the pharmacological profile of a specific leukotriene C(4) binding site not coupled to contraction in human lung parenchyma. Mol Pharmacol 57, 1182–1189 (2000).

46. Ullrich, F., Reincke, S.M., Voss, F.K., Stauber, T. & Jentsch, T.J. Inactivation and Anion Selectivity of Volume-regulated Anion Channels (VRACs) Depend on C-terminal Residues of the First Extracellular Loop. J Biol Chem 291, 17040–17048 (2016).

47. Voets, T., Droogmans, G. & Nilius, B. Modulation of voltage-dependent properties of a swelling-activated Cl^-^ current. Journal of General Physiology 110, 313–325 (1997).

48. Gosling, M., Poyner, D.R. & Smith, J.W. Effects of arachidonic acid upon the volume-sensitive chloride current in rat osteoblast-like (ROS 17/2.8) cells. J Physiol 493 **(Pt** **3**), 613–623 (1996).

49. Rulisek, L. & Vondrasek, J. Coordination geometries of selected transition metal ions (Co^2+^, Ni^2+^, Cu^2+^, Zn^2+^, Cd^2+^, and Hg^2+^) in metalloproteins. J Inorg Biochem 71, 115-127 (1998).

50. Zhou, P., Polovitskaya, M.M. & Jentsch, T.J. LRRC8 N termini influence pore properties and gating of volume-regulated anion channels (VRACs). J Biol Chem 293, 13440–13451 (2018).

51. Deneka, D. et al. Allosteric modulation of LRRC8 channels by targeting their cytoplasmic domains. Nat Commun 12, 5435 (2021).

52. Konig, B., Hao, Y., Schwartz, S., Plested, A.J. & Stauber, T. A FRET sensor of C-terminal movement reveals VRAC activation by plasma membrane DAG signaling rather than ionic strength. eLife 8 (2019).

53. Meyer, K. & Korbmacher, C. Cell swelling activates ATP-dependent voltage-gated Cl^-^ channels in M-1 mouse cortical collecting duct cells. Journal of General Physiology 108, 177–193 (1996).

54. Smart, O.S., Neduvelil, J.G., Wang, X., Wallace, B.A. & Sansom, M.S. HOLE: a program for the analysis of the pore dimensions of ion channel structural models. J Mol Graph 14, 354–360, 376 (1996).

