## Supplementary figures and images for "A conserved mechanism of LRRC8 channel inhibition by distinct drugs"

### Supplemental Figs 1 and 2

#### Supplementary Figure 1

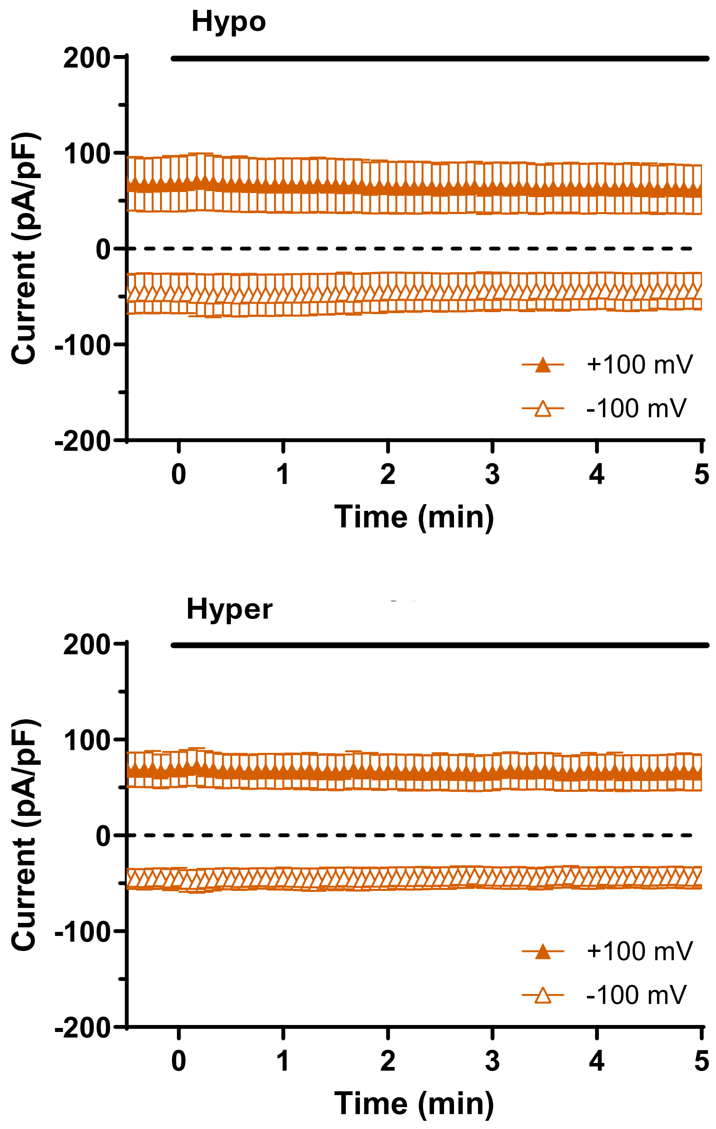

**Supplementary Figure 2**

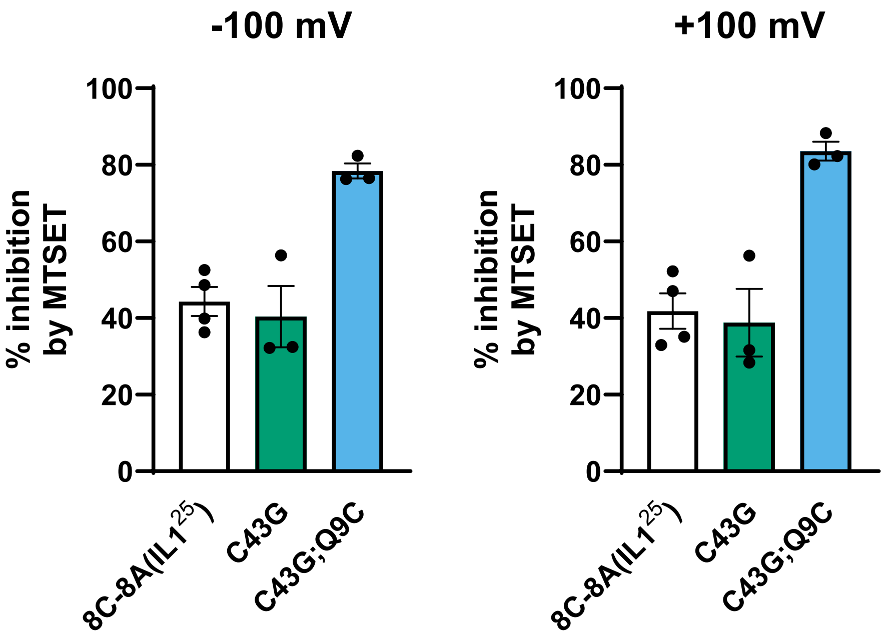
